# Critical roles of Ikaros and HDAC1 in regulation of heterochromatin and tumor suppression in T-cell acute lymphoblastic leukemia

**DOI:** 10.1101/2024.06.27.600861

**Authors:** Yali Ding, Bing He, Daniel Bogush, Joseph Schramm, Chingakham Singh, Katarina Dovat, Julia Randazzo, Diwakar Tukaramrao, Jeremy Hengst, Charyguly Annageldiyev, Avinash Kudva, Dhimant Desai, Arati Sharma, Vladimir S. Spiegelman, Suming Huang, Chi T. Viet, Glenn Dorsam, Giselle Saulnier Scholler, James Broach, Feng Yue, Sinisa Dovat

**Affiliations:** Department of Pediatrics, Pennsylvania State University College of Medicine, Hershey, PA; Department of Medicine, Pennsylvania State University College of Medicine, Hershey, PA; Department of Pharmacology, Pennsylvania State University College of Medicine, Hershey, PA; Department of Oral Maxillofacial Surgery, Loma Linda University School of Dentistry, Loma Linda, CA 92354, USA; Department of Microbiological Sciences, North Dakota State University, Fargo, ND 58102; Department of Biochemistry and Molecular Biology, Pennsylvania State University College of Medicine, Hershey, PA; Northwestern University Feinberg School of Medicine, Chicago, IL

**Author notes:** These authors contributed equally to this work (Yali Ding, Bing He). Corresponding Authors: Sinisa Dovat, M.D., Ph.D., Pennsylvania State University College of Medicine, Department of Pediatrics, 500 University Drive Hershey, PA 17033, USA, Feng Yue, Ph.D., Northwestern University Feinberg School of Medicine, Institute for Artificial Intelligence in Medicine – Center for Advanced Molecular Analysis 675 N. St. Clair, Chicago, IL 60611.

## Abstract

The *IKZF1* gene encodes IKAROS – a DNA binding protein that acts as a tumor suppressor in T-cell acute lymphoblastic leukemia (T-ALL). IKAROS can act as a transcriptional repressor via recruitment of histone deacetylase 1 (HDAC1) and chromatin remodeling, however the mechanisms through which Ikaros exerts its tumor suppressor function via heterochromatin in T-ALL are largely unknown. We studied human and mouse T-ALL using a loss-of-function and *IKZF*1 re-expression approach, along with primary human T-ALL, and normal human and mouse thymocytes to establish the role of Ikaros and HDAC1 in global regulation of facultative heterochromatin and transcriptional repression in T-ALL. Results identified novel Ikaros and HDAC1 functions in T-ALL: Both Ikaros and HDAC1 are essential for EZH2 histone methyltransferase activity and formation of facultative heterochromatin; recruitment of HDAC1 by Ikaros is critical for establishment of H3K27me3 histone modification and repression of active enhancers; and Ikaros-HDAC1 complexes promote formation and expansion of H3K27me3 Large Organized Chromatin lysine (K) domains (LOCKs) and Broad Genic Repression Domains (BGRDs) in T-ALL. Our results establish the central role of Ikaros and HDAC1 in activation of EZH2, global regulation of the facultative heterochromatin landscape, and silencing of active enhancers that regulate oncogene expression.

## Introduction

Ikaros is a transcription factor which acts as a master regulator of hematopoiesis (ref. 1-3) and is essential for immune system development (ref. 4-9) and function (ref. 10,11). Loss of Ikaros function results in high-risk B-ALL (ref. 5,12), T-ALL and early T-cell precursor (ETP) leukemia (ref. 13). Ikaros can activate or repress transcription directly or via recruitment of chromatin remodeling complexes to the promoters of its target genes (ref. 14-26). Several reports demonstrated that Ikaros can regulate gene expression in more complex ways – via global regulation of enhancer landscape (ref. 27), and by regulating global chromatin architecture (ref. 28).

Elucidating the mechanisms through which Ikaros regulates transcription of its target genes is essential for understanding its function as a tumor suppressor in leukemia. A majority of previous reports focused on the role of Ikaros in epigenetic regulation in normal lymphopoiesis and B-ALL, but studies regarding Ikaros function in the epigenomic regulation of transcription in T-ALL are limited (ref. 29-34). Ikaros’ role in transcriptional repression of individual genes has been documented, however its function in global regulation of transcriptional repression in T-ALL has not been studied. Here we use an *IKZF*1 loss-of-function and re-expression approach in human and mouse T-ALL, along with primary human T-ALL, and normal human and mouse thymocytes to determine the roles of Ikaros and HDAC1 in global regulation of facultative heterochromatin and transcriptional repression in T-ALL. Results revealed novel roles of Ikaros and HDAC1 in regulation of EZH2 histone methyltransferase activity, enhancer silencing, and global regulation of formation and expansion of large heterochromatin domains. These data led to a new model of Ikaros function as a global regulator of heterochromatin landscape and gene expression in leukemia.

## Materials and methods

### Cell culture and viral transduction

DN3 cells are Ikaros-null early T-ALL cells that develop spontaneously in Ikaros knock-out mice (ref. 35). DN3 cells were transduced using an MSCV-based bicistronic retroviral vector that expresses Ikaros and GFP. Wild-type cells as well as retrovirally-transduced cells were collected at 1 and 2-day timepoints for experiments.

### RNA-seq

Total RNA was extracted using QIAGEN RNeasy Mini Kit (Qiagen, 74104, Hilden Germany). Libraries were generated and sequenced at the sequencing core facility at Pennsylvania State University College of Medicine (PSUCOM).

### ChIP-seq

ChIP-seq assays for Ikaros, EZH2, HDAC1 and histone modifications were performed using antibodies and methods described in Supplemental Data and as previously described (ref. 36-37). Samples were sequenced using the Illumina HiSeq 2500 at the sequencing core facility at PSUCOM. The rest of experimental procedures and bioinformatics analysis is described in Supplemental Data.

## Results

### Ikaros is essential for formation of facultative heterochromatin

The role of Ikaros in regulation of heterochromatin landscape in T-ALL was studied by comparing Ikaros loss of function and re-expression of Ikaros into Ikaros-null T-ALL cells as previously described **(Fig. S1)** (ref. 27). Ikaros-null T-ALL cells show broad, weak H3K27me3 genome-wide occupancy with very few distinct peaks. Re-expression of Ikaros via retroviral transduction induces abundant H3K27me3 enrichment throughout the genome, predominantly at gene promoters, within 1 day (**Fig. 1A)**. A large majority of newly-enriched H3K27me3 overlaps Ikaros occupancy, suggesting that Ikaros DNA binding directly induces formation of facultative heterochromatin **(Fig. 1B-C**). Motif analysis shows enrichment of Ikaros’ core binding motif in the *de novo*-formed facultative heterochromatin **(Fig. S2)**. Most of the Ikaros-induced *de novo* heterochromatin regions are located within promoters **(Fig. 1C-D)** of the genes that regulate cellular metabolism, signal transduction, and pathways that are essential for malignant transformation, cellular differentiation and proliferation **(Fig. 1E-F).**

**Figure 1.**
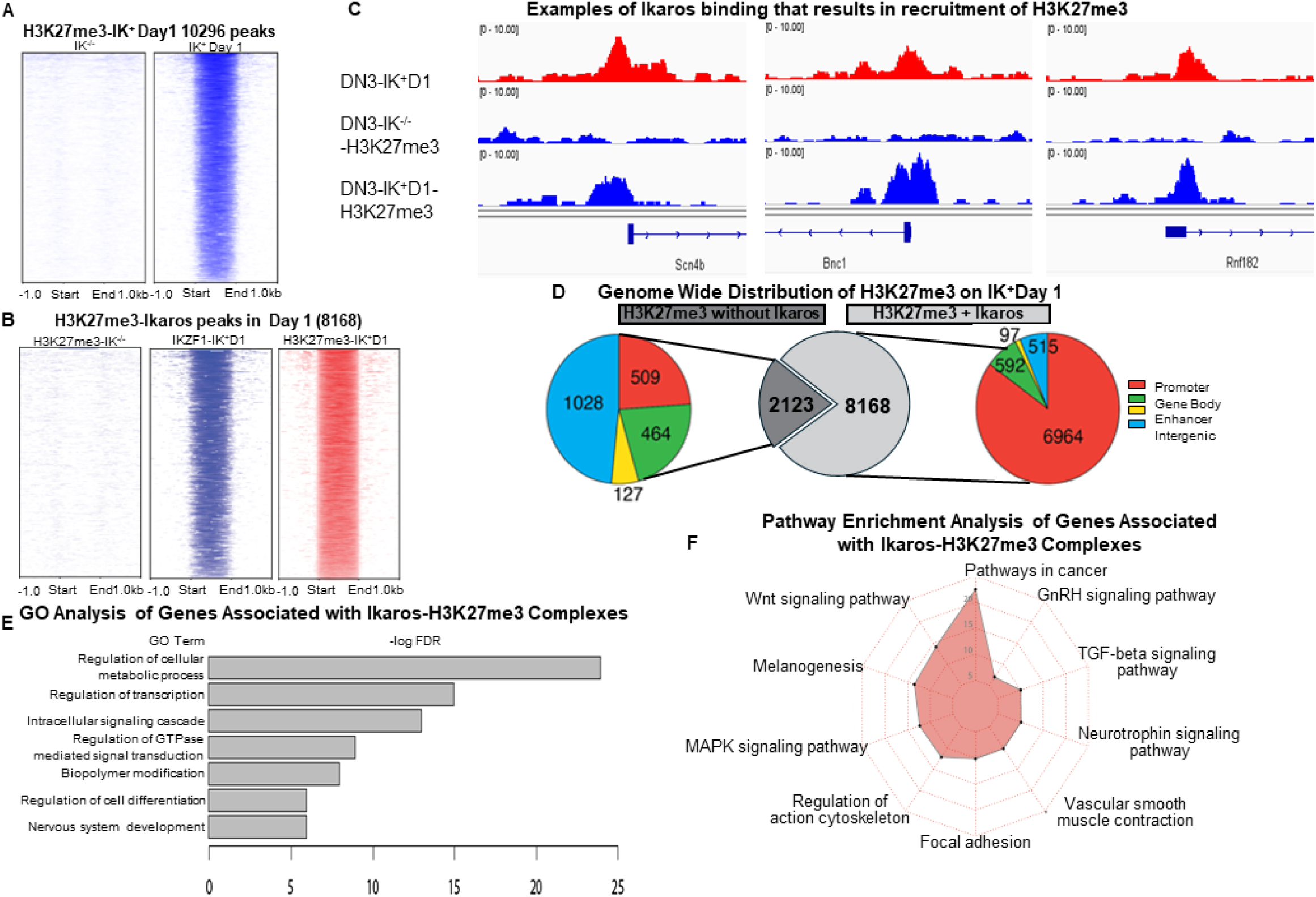
Ikaros binding induces de novo formation of facultative heterochromatin (H3K27me3). (**a**) Heatmaps of H3K27me3 ChIP-seq signals at day 1 after Ikaros introduction vs. day 0. Signals are centered on H3K27me3 peaks at day 1. **(b)** Heatmaps of Ikaros and H3K27me3 ChIP-seq signals with Ikaros direct binding regions at day 0 vs. day 1. **(c)** Examples of *de novo*-formed H3K27me3 enrichment that are induced by Ikaros binding. Ikaros-induced epigenetic changes are shaded in grey. **(d)** *De novo* H3K27me3 regions classified by function of the DNA element. **(e-f)** Gene ontology and pathway enrichment analysis of genes associated with *de novo* H3K27me3 regions.

Analysis of H3K27me3 enrichment in the subsequent day following Ikaros-re-expression revealed a dual effect of Ikaros on the facultative heterochromatin landscape: 1) direct formation of facultative heterochromatin is associated with Ikaros occupancy and occurs mostly at the promoters of target genes and 2) establishment of global facultative heterochromatin landscape without Ikaros occupancy (indirect effect). Direct formation of facultative heterochromatin by Ikaros is predominant the first day following Ikaros re-expression. In contrast, the global increase (over 4-fold) of H3K27me3 peaks in facultative heterochromatin occurs mostly independent of concomitant Ikaros DNA binding at the 2^nd^ day following Ikaros re-expression and is similarly distributed among promoters, gene body and intergenic regions **(Fig. S3).** Dynamic changes in the H3K27me3 signature during the first 2 days following Ikaros re-expression is consistent with establishment of the heterochromatin landscape that is found in normal thymocytes **(Figs. S4-S5).** These data suggest that Ikaros is essential for formation of facultative heterochromatin and that Ikaros regulates H3K27me3 landscape both by direct DNA binding, and by indirectly, likely by influencing the activity of genes that regulate H3K27me3 formation. Genome-wide distribution of H3K27me3 following Ikaros re-expression suggests Ikaros-driven heterochromatin re-programming to that of a physiological thymocyte.

### Interaction between EZH2 and Ikaros is required for formation of facultative heterochromatin

Enhancer of zeste homolog 2 (EZH2) is a histone methyltransferase that catalyzes the trimethylation of Histone 3 at lysine 27 resulting in H3K27me3 repressive chromatin (ref. 38-39). Ikaros can bind and recruit EZH2 to promoters of Ikaros target genes resulting in the formation of facultative heterochromatin (ref. 40). We analyzed how the absence of Ikaros and its re-expression affect function of EZH2. ChIP-seq showed abundant EZH2 occupancy in Ikaros-null T-ALL, mostly at promoter regions, but without enrichment of H3K27me3 **(Fig. 2A, S6A).** Re-expression of Ikaros, results in strong recruitment of EZH2 predominantly by Ikaros, and was associated with H3K27me3 enrichment **(Fig. 2B-left heat maps, S6B)**. DNA-bound EZH2 not associated with Ikaros showed poor enrichment in H3K27me3 **(Fig. 2C, S6C).** Interestingly, many of the sites occupied solely by EZH2 in day 0 without H3K27me3 enrichment, were occupied by Ikaros-EZH2 complexes in day #1 and resulted in H3K27me3 enrichment **(Fig. S6D).** The relationship between EZH2 and Ikaros peaks in IK-null T-ALL and 1 day following Ikaros re-expression is shown in **Figure S7**. Results in **Figure 2A-C** suggest that EZH2 DNA occupancy without concomitant Ikaros DNA binding is not sufficient to induce H3K27me3 enrichment, and that EZH2 requires Ikaros binding for the formation of facultative heterochromatin. Ikaros-EZH2 complexes are mostly located at the promoters of their target genes **(Fig. 2D),** which regulate protein metabolism and signaling pathways in cancer (Wnt, MAPK, etc.) **(Fig. 2E-F).**

**Figure 2.**
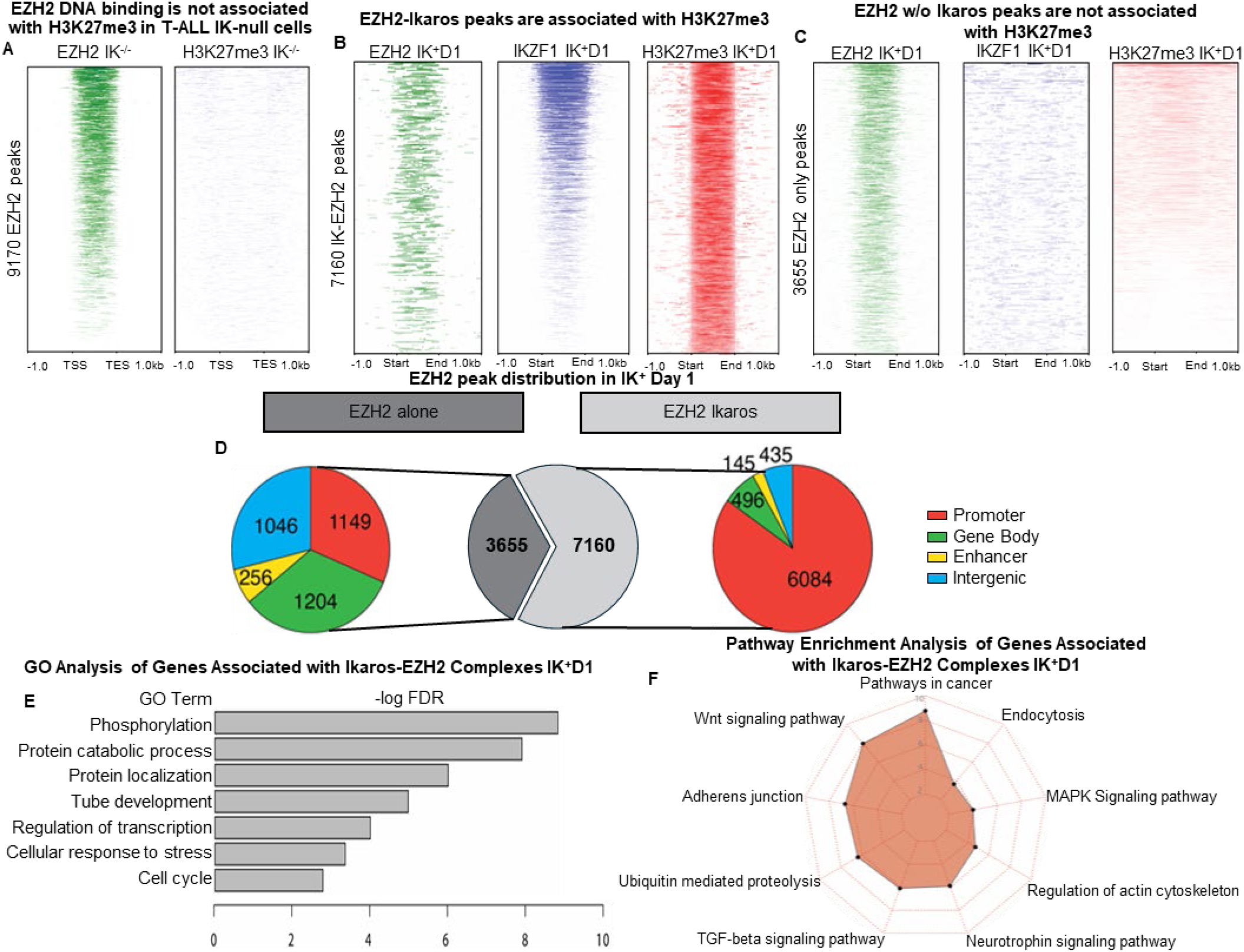
Ikaros recruits and activates EZH2. (**a**) Heatmaps of EZH2 and H3K27me3 ChIP-seq signals in Ikaros-null T-ALL. **(b-c)** Heatmaps of EZH2, Ikaros and H3K27me3 ChIP-seq signals in Ikaros-EZH2 occupied **(b),** and EZH2-only occupied **(c)** regions. **(d)** EZH2-only and EZH2-Ikaros occupied sites classified by function of the DNA element. **(e-f)** Gene ontology and pathway enrichment analysis of genes associated with EZH2-Ikaros complexes.

Analysis of EZH2 enrichment in the subsequent day showed reduced occupancy of EZH2 and Ikaros-EZH2 complexes, especially at the promoters of target genes **(Fig. S8).** Overall, these data suggest that formation of Ikaros-EZH2 complex is essential for H3K27me3 enrichment and formation of facultative heterochromatin the first day after Ikaros re-expression, and that EZH2 DNA binding in Ikaros-null T-ALL is not associated with H3K27me3 enrichment.

### Ikaros recruits HDAC1 to the promoters of the genes that regulate malignant transformation

Ikaros can recruit HDAC1 to regulate expression of its target genes via chromatin remodeling (ref. 14-15, 22, 41-42). We analyzed whether the lack of Ikaros and its re-expression regulate HDAC1 genome-wide distribution and function. In Ikaros-null T-ALL, HDAC1 DNA occupancy is poor and occurs mostly at intergenic regions (**Fig. 3A and S9-top panel**). In day 1 following Ikaros re-expression, there is a strongly increased HDAC1 DNA occupancy due to 1) recruitment of HDAC1 by Ikaros, mostly to promoters **(Fig. 3B-C)**; and 2) due to global genomic occupancy of HDAC1 that is independent of Ikaros binding **(Fig. 3B, S9)**. Ikaros-HDAC1 complexes localize to promoters and are associated mostly with repression of their target genes that regulate protein metabolism and oncogenic signaling pathways **(Fig. 3D-F)**, similarly to the pathways regulated by EZH2 and H3K27me3.

**Figure 3.**
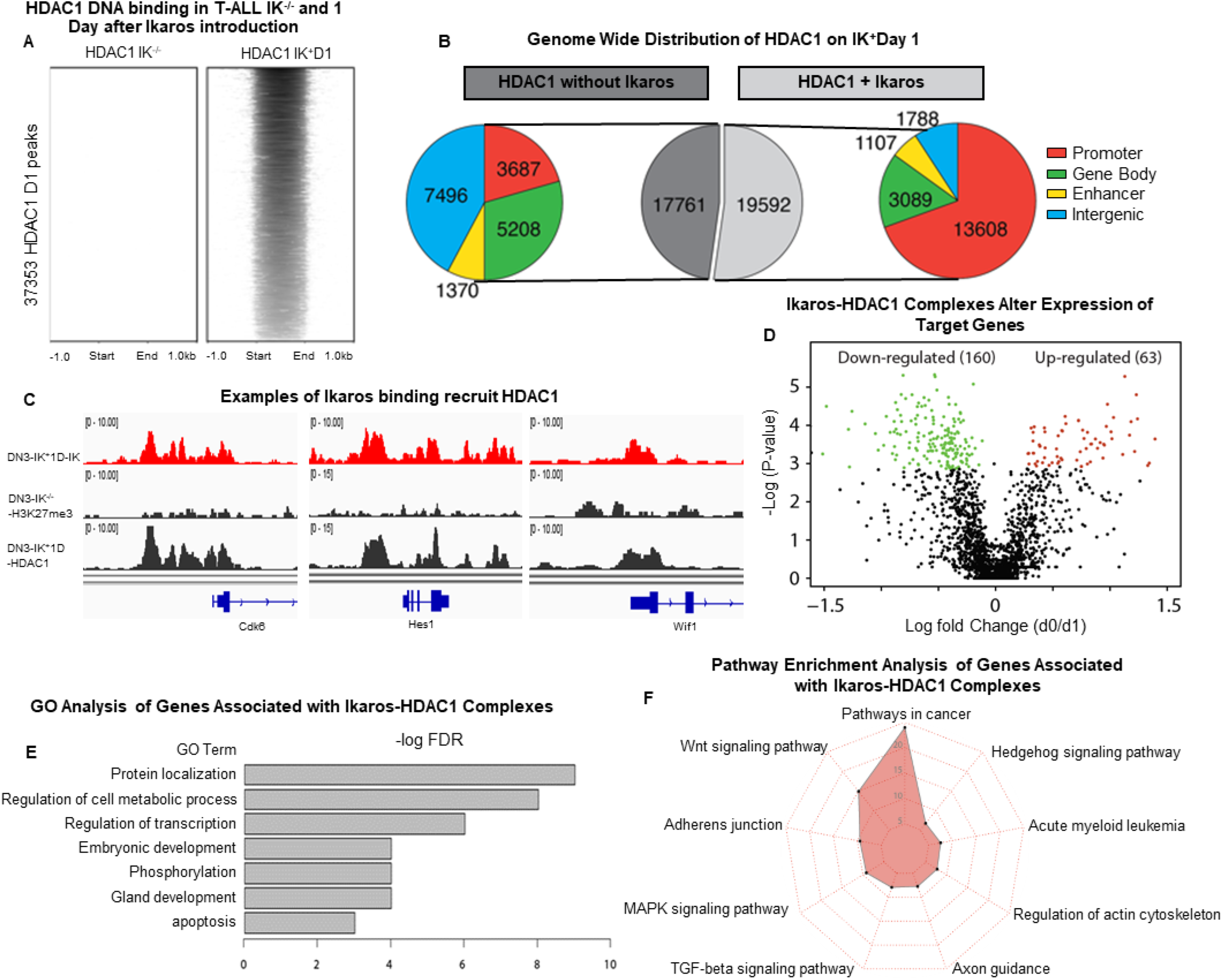
Ikaros is essential for HDAC1 recruitment to the gene promoters. (**a**) Heatmaps of HDAC1 ChIP-seq signals in IK-null T-ALL and day 1 following Ikaros re-expression. **(b)** HDAC1-only and HDAC1-Ikaros occupied sites classified by function of the DNA element. **(c)** Examples of Ikaros recruitment of HDAC1 to the promoters of Ikaros target genes. **(d)** Analysis of the differentially expressed genes directly regulated by Ikaros-HDAC1 complexes. **(e-f)** Gene ontology and pathway enrichment analysis of genes associated with Ikaros-HDAC1 complexes.

Analysis of HDAC1 occupancy in second day following Ikaros re-expression, showed highly increased HDAC1 genome-wide occupancy, with Ikaros-HDAC1 complexes present mostly at the promoters of Ikaros target genes. However, most HDAC1 global DNA enrichment was not associated with concomitant Ikaros binding, and is similarly distributed among promoters, gene body and intergenic regions **(Fig. S9).**

### HDAC1 has a critical role in formation of facultative heterochromatin

We further analyzed the relationship between Ikaros, EZH2 and HDAC1 with H3K27me3 enrichment and formation of facultative heterochromatin. Surprisingly, H3K27me3 enrichment has the strongest association with HDAC1 DNA occupancy, much stronger than with EZH2 and/or Ikaros occupancy **(Fig. 4. A-B)** suggesting that HDAC1 has a more prominent role in formation of H3K27me3 than EZH2. The role of HDAC1 in H3K27me3 formation is even more prominent at promoter regions **(Fig. S10).** Dynamic analysis of Ikaros, EZH2, and HDAC1 occupancy with H3K27me3 showed that initial formation of H3K27me3 (day 1) is associated either with Ikaros/HDAC1/EZH2 concomitant DNA binding, or with Ikaros/HDAC1 DNA binding without EZH2 occupancy **(Fig. 4A).** However, further H3K27me3 enrichment (day 2) is either associated with HDAC1 occupancy independent of Ikaros and/or EZH2 enrichment, or independent of Ikaros/HDAC1/EZH2 DNA binding altogether **(Fig. 4B).** These results strongly suggest that the recruitment of HDAC1 by Ikaros is essential for induction of H3K27me3 and that HDAC1 is critical for the maintenance of H3K27me3.

**Figure 4.**
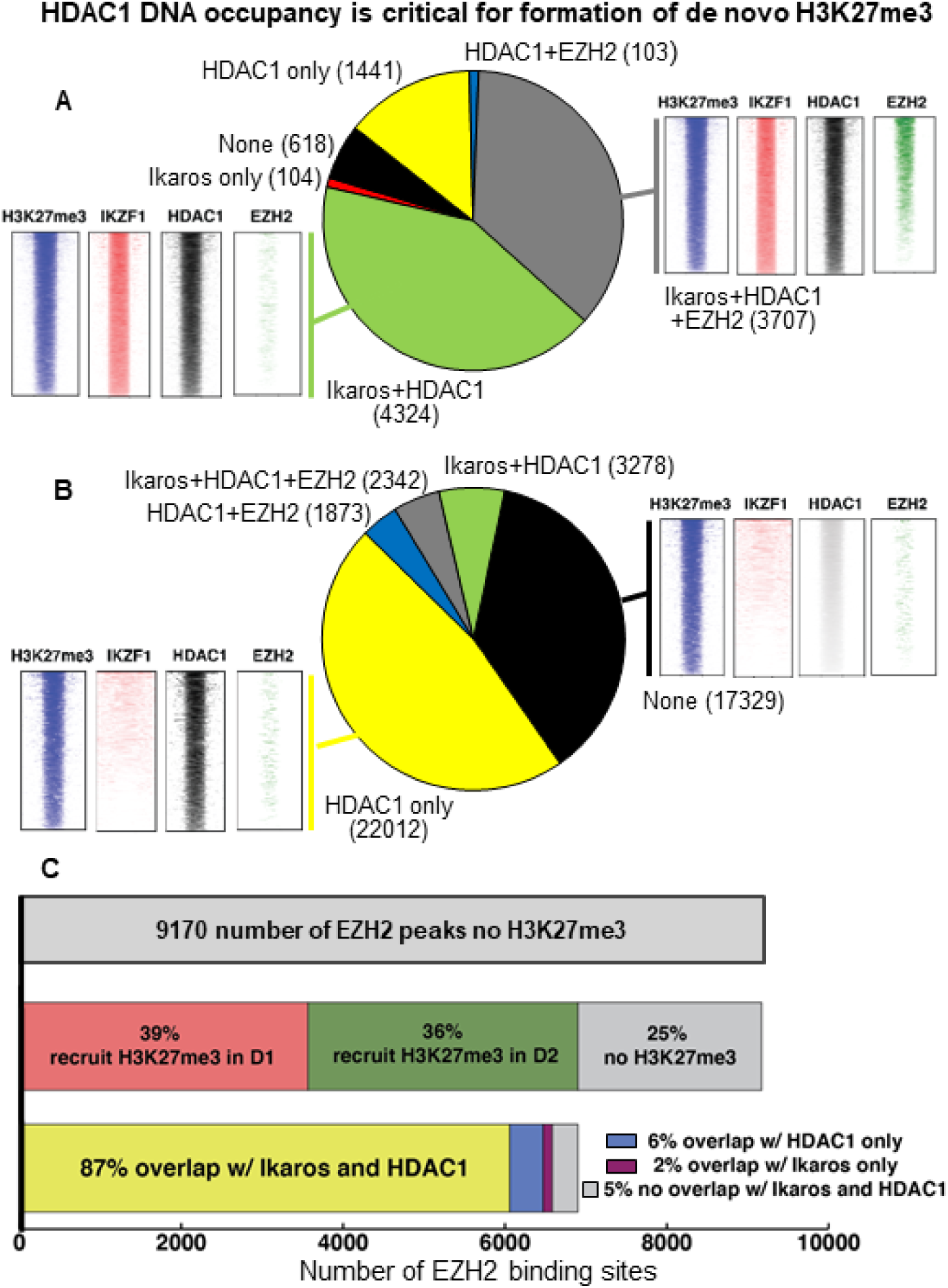
Critical roles of Ikaros and HDAC1 in the global de novo formation of H3K27me3. (**a-b**) Ikaros, HDAC1, and EZH2 occupancy associated with H3K27me3 formation in days 1-2 following Ikaros re-expression. **(c)** EZH2 target genes are regulated by the Ikaros and HDAC1-induced epigenetic switch. EZH2 DNA occupancy at target genes does not induce H3K27me3 in Ikaros-null T-ALL (top). A large number (75%) of EZH2-target genes undergo epigenetic switch with de novo H3K27me3 formation following Ikaros-re-expression (middle). Most of the EZH2 target genes that undergo epigenetic switch are regulated by Ikaros and HDAC1 binding (bottom).

The role of HDAC1 recruitment by Ikaros, as well as global genomic occupancy of HDAC1 is highly prominent during the first 2 days following Ikaros re-expression and during proliferation arrest of Ikaros-null T-ALL cells. Since EZH2 DNA binding did not induce formation of H3K27me3 in Ikaros-null cells, we analyzed whether re-expression of Ikaros/HDAC1 induces H3K27me3 enrichment at sites that were occupied by EZH2 in Ikaros-null T-ALL. Results showed that Ikaros re-expression induced H3K27me3 enrichment at 75% of the sites occupied by EZH2 in Ikaros-null cells **(Fig. 4C-middle).** Almost all of the H3K27me3-enriched sites were occupied by both Ikaros and HDAC1 **(Fig. 4C-bottom)** and most of them were located at promoters of the genes that regulate oncogenic signaling pathways **(Fig. S11).** These results suggest that recruitment of HDAC1 by Ikaros is essential for EZH2 tumor suppressor function by inducing formation of facultative heterochromatin at promoters of the genes that positively regulate oncogenic pathways.

### Ikaros and HDAC1 modulate activity of enhancers

Next, we analyzed how recruitment of HDAC1 by Ikaros regulates enhancer activity. A large set of enhancers are occupied by either Ikaros or HDAC1 during the first two days following Ikaros re-expression into Ikaros-null cells **(Fig. 5A).** We compared transcription of genes that are regulated by the active enhancers (H3K4me1+/H3K27ac+), which are not occupied by Ikaros, nor HDAC1 (Ikaros-/HDAC1-) to the transcription of the genes regulated by enhancers that are enriched for Ikaros (Ikaros+), HDAC1 (HDAC1+) or both Ikaros and HDAC1 (Ikaros+/HDAC1+). Ikaros occupancy at active enhancers has a repressive effect on transcription of target genes of those enhancers **(Fig. 5B)**. HDAC1 enrichment has an even more pronounced repressive effect on enhancer activity, similar to concomitant Ikaros and HDAC1 enrichment **(Fig. 5B).** Modulation of enhancer function by Ikaros and HDAC1, represses transcription of the genes that regulate cancer pathways, (e.g. hippo, Ras, etc.) **(Fig. S12-14)**.

**Figure 5.**
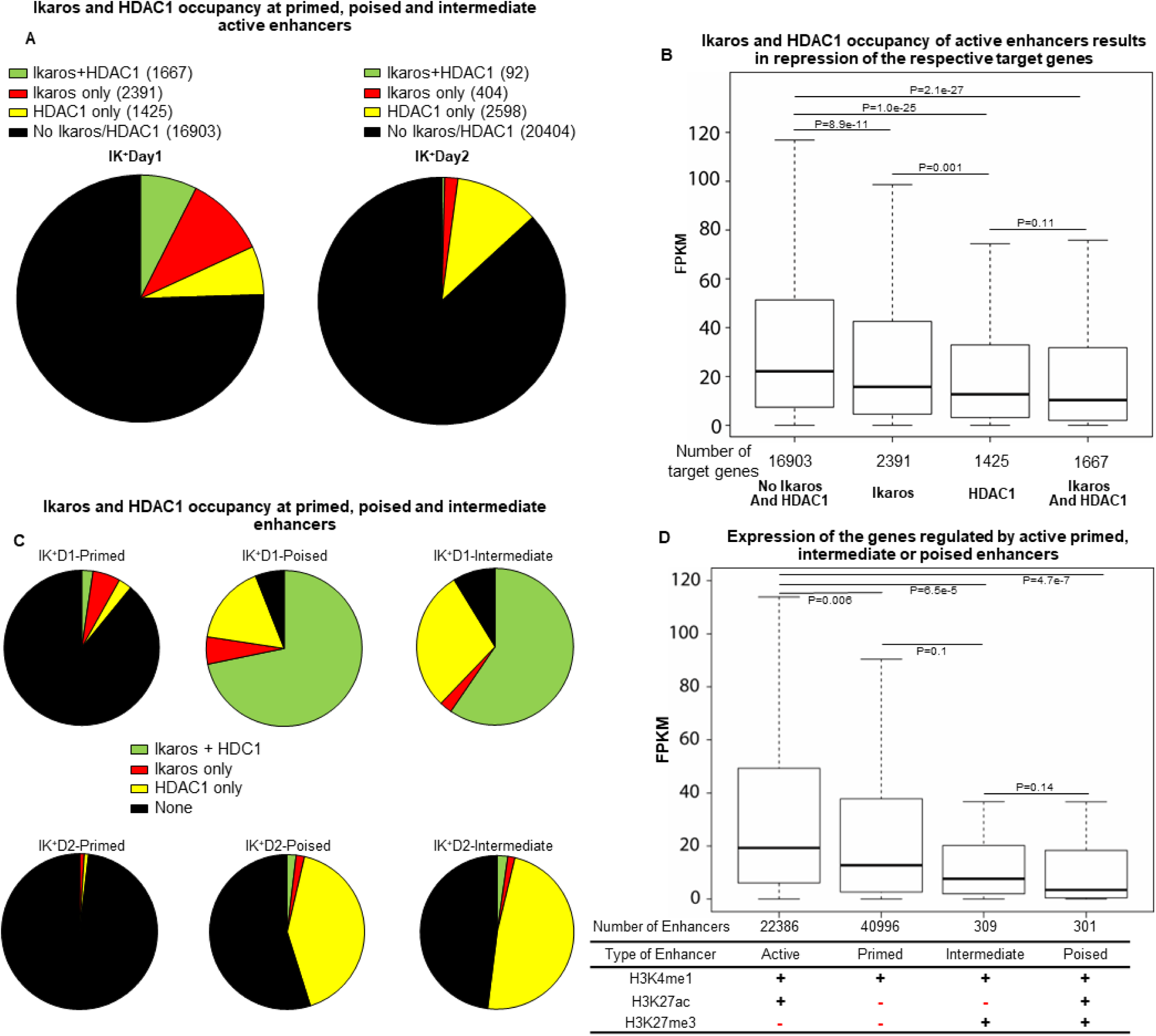
Ikaros and HDAC1 are repressors of active enhancers. (**a**) A large number of active enhancers are occupied by Ikaros and HDAC1 after Ikaros re-expression. Pie charts showing the portions of active enhancers occupied by Ikaros, HDAC1, or both Ikaros and HDAC1 **(b)** Boxplot shows expression of genes regulated by the active enhancers not bound by Ikaros and/or HDAC1 (left) and occupied by Ikaros and/or HDAC1. **(c)** Ikaros and HDAC1 regulate formation of poised and intermediate enhancers. Pie charts showing the portions of primed, poised, and silenced enhancers occupied by Ikaros, HDAC1, or both Ikaros and HDAC1 in day 1 and 2 following Ikaros re-expression. Ikaros-HDAC1 co-occupancy is highly associated with formation of poised and intermediate enhancers in day 1, while HDAC1-only occupancy is dominant in those enhancers in day 2. **(d)** Boxplot shows expression of genes regulated by the active, primed, intermediate, and poised enhancers.

Non-active enhancers can be classified as primed (enriched in H3K4me1+ only), intermediate (H3K4me1+/H3K27ac+/H3K27me3+) and poised (H3K4me1+/H3K27me3+) (ref. 43-44). We analyzed whether Ikaros and HDAC1 can regulate formation of these enhancer types. Results show that a majority of poised and intermediate enhancers in day 1 following Ikaros re-expression are associated with both Ikaros/HDAC1 occupancy, while HDAC1 occupancy has a dominant role in formation of poised and intermediate enhancers during the second day following Ikaros re-expression into Ikaros-null T-ALL **(Fig. 5C)**. Genes regulated by poised and intermediate enhancers have significantly reduced transcription compared to the genes regulated by active or primed enhancers **(Fig. 5D).** These data show that Ikaros and HDAC1 occupancy can silence enhancer activity in two ways: 1) by repressing activity of active enhancers; and 2) by inducing formation of poised and/or silenced enhancers. To study the functional significance of Ikaros and HDAC1 regulation of enhancer activity in T-ALL, we analyzed gene regulation by active enhancers in Ikaros-null T-ALL that become occupied by Ikaros and/or HDAC1 following Ikaros transduction **(Fig. S15**). Pathway enrichment analysis show that these enhancers regulate genes involved in pathways in cancers and cell proliferation (**Fig. S15A**). Ikaros re-expression and Ikaros/HDAC1 binding to those enhancers results in repression of many genes that promote stemness and drug resistance, e.g. adrenomedullin **(Fig. S15B)**. These data suggest that Ikaros exerts its tumor suppressor function in T-ALL by recruiting HDAC1 and silencing activity of enhancers that regulate transcription of various oncogenes. Results identified enhancers and their corresponding pathways that are directly regulated by Ikaros in T-ALL.

### Ikaros regulates formation and expansion of Large Organized Chromatin lysine (K) domains (LOCKs)

Global epigenetic landscape contains clusters of nucleosomes with a large number of post-transcriptionally modified lysine residues. These clusters define Large Organized Chromatin lysine (K) domains (LOCKs) which are associated with repression if they consist predominantly of heterochromatin marks (H3K27me3 or H3K9me3) (ref. 45). We analyzed whether Ikaros can regulate formation and distribution of LOCKs. Ikaros expression strongly induces de novo formation of H3K27me3 LOCKs, which becomes even more prominent 2 days following Ikaros re-expression **(Fig. 6A).** Re-expression of Ikaros increases the number of H3K27me3 LOCKs but has a greater effect by increasing the genomic coverage of H3K27me3 LOCKs **(Fig. 6A).** This is mostly due to strong day 2 expansion of the de novo formed H3K27me3 LOCKs in day 1 **(Fig. 6B).** Expansion of the H3K27me3 LOCKs over 2 days following Ikaros re-expression results in a large number of genes being regulated by LOCKs. **(Fig. 6C).** There is a dynamic shift in the function of the genes regulated by H3K27me3 LOCKs: in day #1 following Ikaros re-expression, H3K27me3 LOCKs regulate mostly pathways in cancer and cellular proliferation, while in day #2, H3K27me3 LOCKs mostly regulate genes involved in immune response **(Fig. S16)**. Expression of genes regulated by H3K27me3 LOCKs is reduced compared to the expression of the random genes **(Fig. 6D)**. These data demonstrate that Ikaros has an important role in organization of higher heterochromatin structures and gene expression via regulation of formation and expansion of H3K27me3 LOCKs.

**Figure 6.**
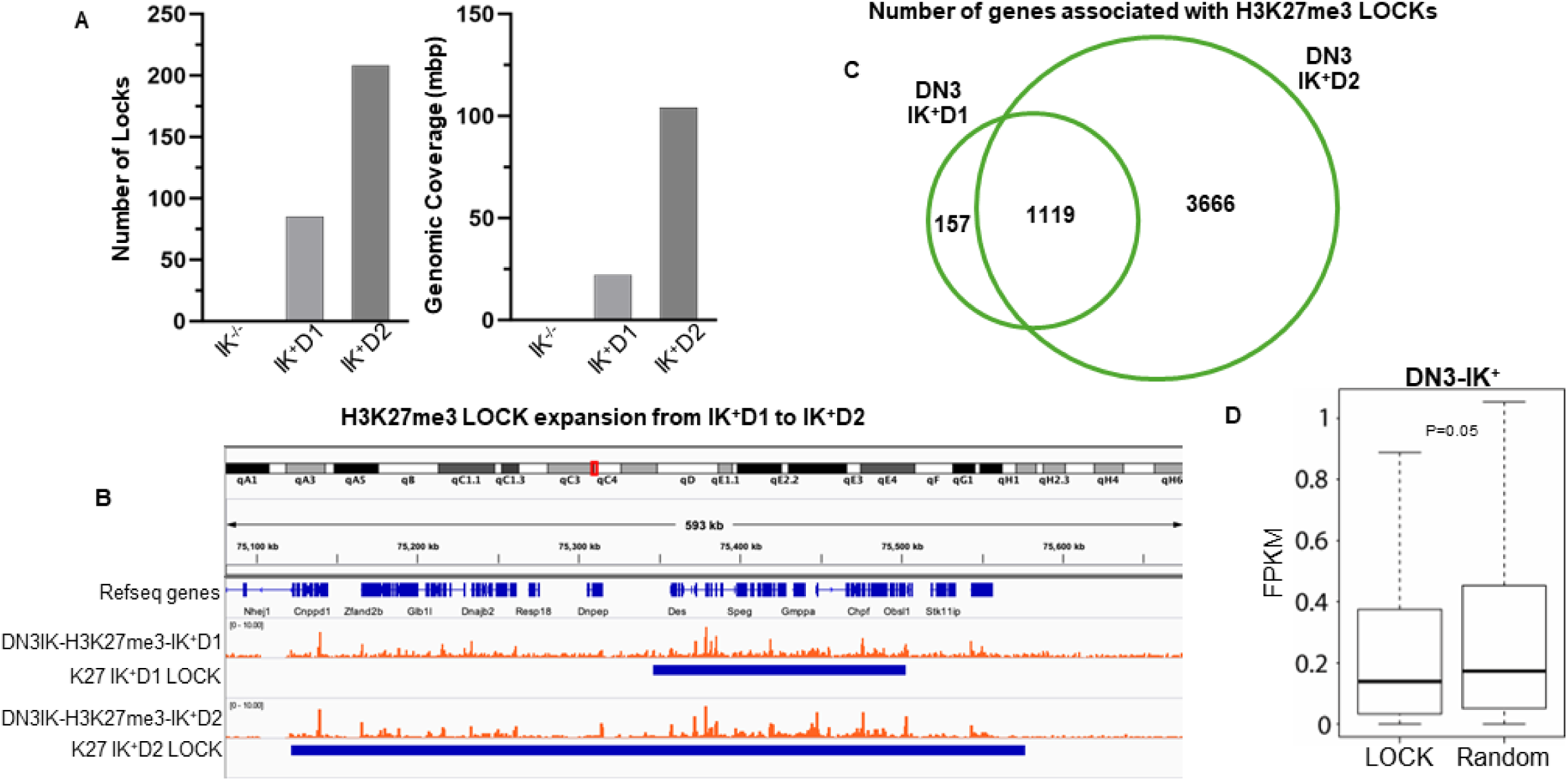
Ikaros regulates formation and expansion of H3K27m3 LOCKs. (**A**) Number (left) and genomic coverage (right) of H3K27me3 LOCKs in day 1 and day 2 following Ikaros re-expression. **(B)** Example of the genomic expansion of the H3K27me3 LOCKs from day 1 to day 2 after Ikaros re-expression. **(C)** Number of genes regulated by the H3K27me3 LOCKs in day 1 and day 2 following Ikaros re-expression. **(D)** Gene expression of the genes regulated by LOCKs vs. random genes.

### Regulation of heterochromatin landscape by Ikaros in T-ALL is evolutionarily conserved

Ikaros functions as a tumor suppressor in human T-ALL (ref. 13). We tested whether Ikaros regulates heterochromatin landscape in the same way in mouse and human T-ALL. The IKZF1 gene was deleted in human T-ALL cells MOLT-4 using the CRISPR-Cas system to create MOLT-4 IKZF1-null cells. We analyzed Ikaros’ role in heterochromatin regulation in human T-ALL by comparing the heterochromatin landscape of MOLT-4-IKZF1-null and MOLT-4 wild type cells. IKZF1 deletion in MOLT-4 cells resulted in reduced global H3K27me3 and HDAC1 occupancy **(Fig. 7A).** This was associated with the global genomic redistribution of H3K27me3 and HDAC1 due to their reduced occupancy at promoter regions **(Fig. 7B, Fig. S17)**. Ikaros knockout reduced the H3K27me3 and HDAC1 occupancy at promoters of genes that regulate cell cycle progression, cellular division, and nucleotide metabolism **(Fig. 7C-D).** Gene expression analysis showed reduced expression of the genes regulated by active enhancers with Ikaros and HDAC1 occupancy compared to the expression of the genes regulated by the active enhancers not occupied by Ikaros and HDAC1 in human T-ALL **(Fig. S18).** Ikaros deletion in MOLT-4 cells reduced both the number and the genomic coverage of H3K27me3 LOCKS **(Fig. S19).** Lack of Ikaros reduces the number of genes that are regulated by LOCKs, including genes involved in several oncogenic pathways **(Fig. S20).** We analyzed the effect of Ikaros deletion on formation of the Broad Genic Repression Domains (BGRD) in MOLT-4 cells. BGRDs are domains enriched in H3K27me3 **(Fig. S21A)** that are also enriched in oncogenes (ref. 46). Expression of genes located within BGRDs is strongly reduced compared to that of random genes **(Fig. S21B)**. Ikaros deletion reduces the number of BGRDs and its genomic coverage **(Fig. S21C-D).** We analyzed the H3K27me3 ChIP-seq data from mouse Ikaros-null T-ALL on day 1 and 2 following Ikaros re-expression and human primary T-ALL, and then compared them with the published H3K27me3 genomic occupancy in normal mouse and human thymocytes. Comparative analysis of genome wide enrichment of H3K27me3 in normal and malignant human and mouse cells showed a remarkable conservation of H3K27me3 landscape in human and mouse thymocytes, loss of H3K27me3 in T-ALL, and an important role of Ikaros in maintaining H3K27me3 landscape **(Fig. S22).** These data suggest that Ikaros regulates global heterochromatin landscape in a similar way for both mouse and human T-ALL, validating the mouse T-ALL model used for this study and providing insight into the tumor suppressor role of IKAROS in human T-ALL.

**Figure 7.**
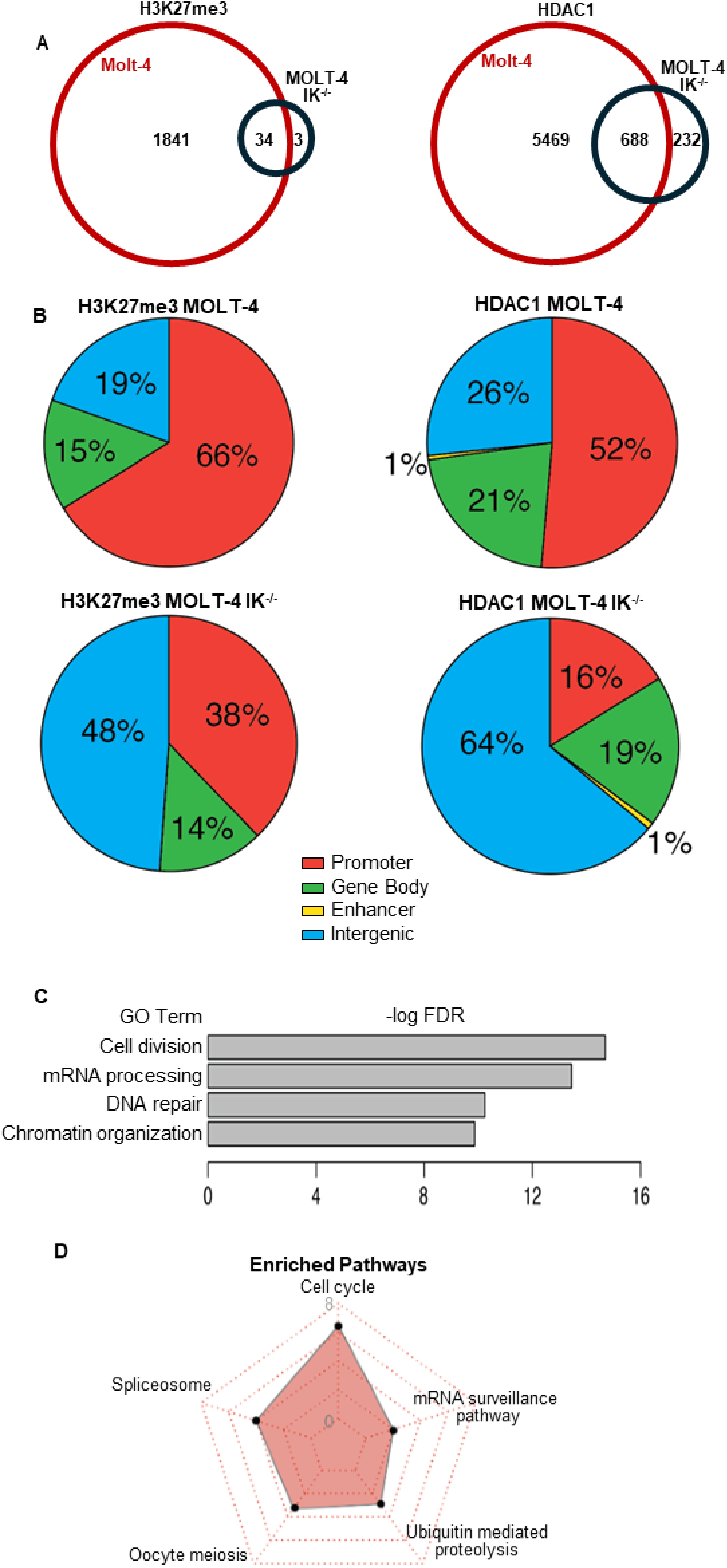
Ikaros regulates genomic occupancy (A) and distribution (B) of EZH2 and HDAC1 and H3K27me3 formation in human T-ALL. (**A**) Number of H3K27me3 (left) and HDAC1 (right) peaks in wildtype MOLT-4 and in Ikaros-knockout (MOLT-IK-KO) cells. **(B)**, H3K27me3 and HDAC1 occupied sites, in wildtype MOLT-4 and in Ikaros-knockout (MOLT-IK-KO) cells, classified by function of the DNA element. **(C-D)** Gene ontology and pathway enrichment analysis of genes associated with HDAC1 occupancy in wildtype MOLT-4 cells.

**Figure 8.**
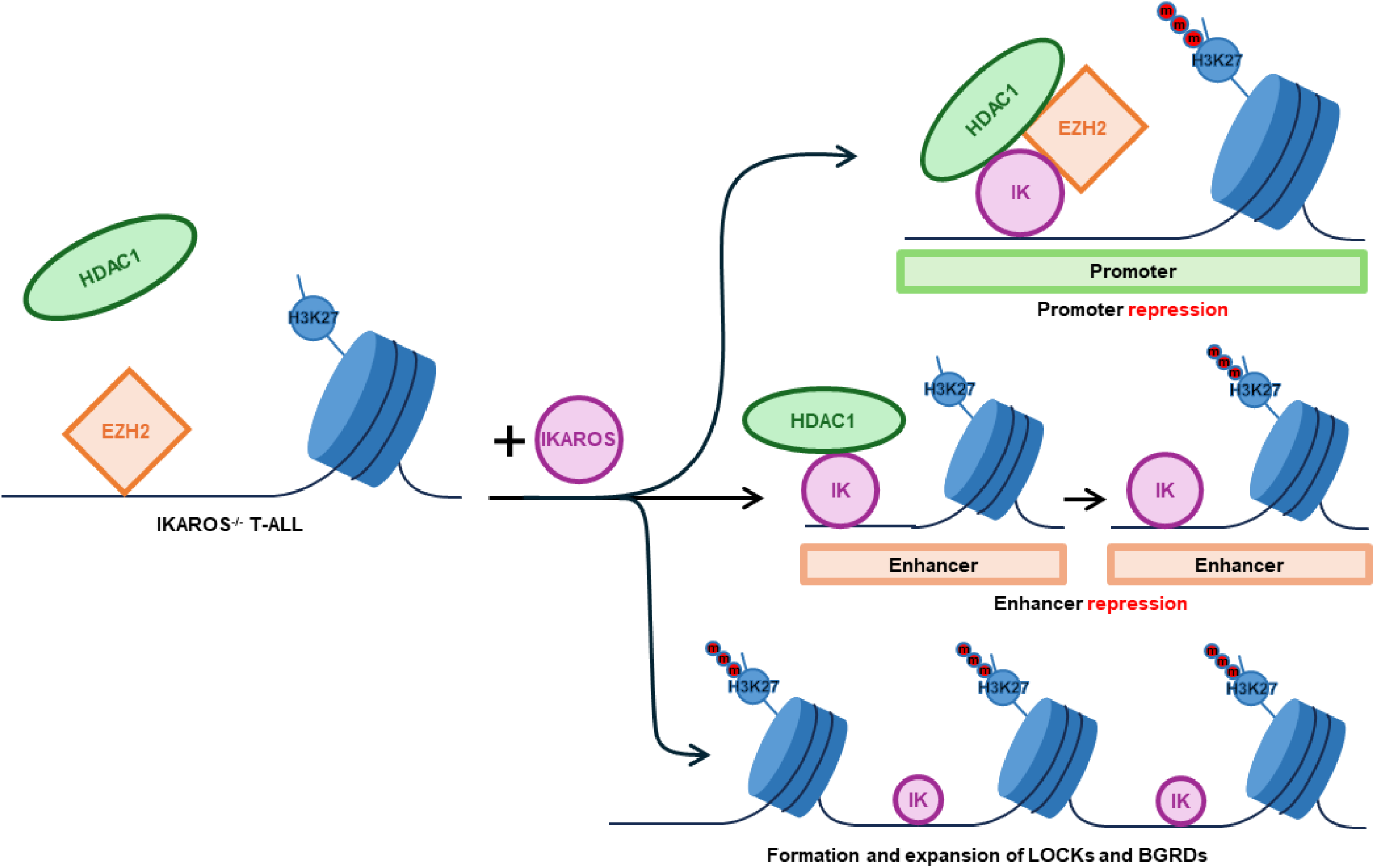
Novel Ikaros functions in the regulation of heterochromatin formation. Ikaros expression regulates: *de novo* formation of H3K27me3 by HDAC1 recruitment and activation of EZH2; repression of active enhancers; and formation and expansion of LOCKs and BGRDs.

## Discussion

Ikaros is a tumor suppressor in T-ALL. Ikaros loss-of-function and re-expression experiments in human and mouse T-ALL, coupled with extensive epitranscriptomic analysis of primary T-ALL and thymocytes, identified novel mechanisms that regulate tumor suppression in T-ALL:

a. *Critical role of Ikaros-HDAC1 complexes in regulation of global facultative heterochromatin landscape.* Loss of the H3K27me3-associated heterochromatin has been associated with development of T-ALL and other types of leukemia (ref. 13, 47-52). Inactivating mutations of histone methyltransferase EZH2 is considered the principal mechanism responsible for the loss of H3K27me3 in T-ALL (ref. 53-56). Presented data revealed that DNA binding of EZH2 in the absence of Ikaros and/or HDAC1 co-localization is rarely associated with H3K27me3 in both Ikaros-null and Ikaros-wildtype mouse and human T-ALL. Lack of Ikaros results in severe depletion of DNA-bound HDAC1, and loss of EZH2-associated H3K27me3. Thus, HDAC1 recruitment by Ikaros is critical for both EZH2 activity and H3K27me3 formation.
b. *Central role of HDAC1 in EZH2 activation, H3K27me3 formation and maintenance*. Results showed that the recruitment of HDAC1 by Ikaros is essential for EZH2 function and de novo H3K27me3 formation. Since Ikaros recruits both HDAC1 and EZH2, and HDAC1 can interact with EZH2 (ref. 57), it is possible that HDAC1 directly activates EZH2 while in complex with Ikaros. HDAC1 DNA occupancy does not result in loss of H3K27ac, but rather in de novo formation and genome-wide expansion of H3K27me3.
c. *Ikaros/HDAC1 complexes can repress active enhancers and promote formation of intermediary and poised enhancers*. Ikaros can induce de novo formation of enhancers and activation of primed enhancers (ref. 27). Presented data showed that Ikaros binding is associated with silencing of enhancers’ activity. This can occur via recruitment of HDAC1 without H3K27me3 formation (**Fig. 5B**), or via formation of intermediate or poised enhancers **(Fig. 5D).** These results, together with previously reported data, reveal a complex regulatory role of Ikaros in enhancer activity: Ikaros DNA binding can result in de novo formation and/or activation of enhancers, but also in repression of enhancers’ activity. Ikaros-silenced enhancers regulate oncogenic pathways in Ikaros-null T-ALL **(Fig. S15),** which suggests that silencing enhancers that regulate expression of oncogenes is one of the mechanisms through which Ikaros exerts a tumor suppressor function in T-ALL.
d. *Regulation of formation and expansion of LOCKs and BGRDs*. Lack of Ikaros results in severe depletion of H3K27me3 LOCKs and BGRDs. These recently identified large heterochromatin domains (BGRDs) strongly repress a large sets of genes, including oncogenes, and have a prominent role in global epigenomic regulation of gene expression (ref. 45-46). Ikaros-driven HDAC1 recruitment and H3K27me3 expansion is a likely mechanism through which Ikaros promotes formation of LOCKs and BGRDs. Results suggest that Ikaros-HDAC1 complexes could function as a tipping point at the intersection between euchromatin and heterochromatin in T-ALL. Ikaros’ prominent roles in both formation of super-enhancers and large heterochromatin domains, suggest that Ikaros regulates higher chromatin organization.

In conclusion, our results identify novel functions of Ikaros and HDAC1 in regulation of heterochromatin propagation and enhancer activity. The presented data demonstrate that Ikaros-HDAC1 complexes regulate the balance between euchromatin and heterochromatin in T-ALL. Results reveal the novel mechanisms that regulate interplay between active and repressive chromatin, global regulation of gene expression, and tumor suppression in leukemia.

## Acknowledgements

This work was supported by R01CA209829, R01CA213912, and (SD); UM1HG012649; R35GM124820 (FY); R01HG011207 (FY and SD), T32CA060395 (JS), Hyundai Hope on Wheels Scholar Grant (to SD and JS), and the Four Diamonds Fund of the Pennsylvania State University College of Medicine (to SD and JS).

## Supplemental Methods

### Cell culture

Molt4 Ikaros CRISPR knockout and control cells were purchased form Synthego, Redwood City, CA. The sequence of sgRNA for knockout Ikaros gene is AUCUGGAGUAUCGCUUACAG. Cells were cultured with RPMI 160 with 10% FBS. Both PCR and western blot were performed to confirm the knockout of Ikaros. Human primary T-ALL PDX cells W8 were obtained from Loma Linda University (Loma Linda, CA) in compliance with Institutional Review Board regulations.

### Western blot

10^6^ cells were collected and lysed by Pierce IP Lysis Buffer (Pierce, 87787) on ice for 10 minutes. Cell debris were removed by centrifugation at 13, 000g at 4℃ for 10 minutes. Supernatants were transferred to new tubes for Western blot which was performed according to standard procedure. Antibodies which were used include anti-HA tag (Abcam, ab9110) for HA-tagged Ikaros, anti-HDAC1 (Abcam, ab7028), anti-EZH2 (Active Motif, 39901) and anti-PCNA (Santa Cruz, sc-7907).

### ChIP-seq

Antibodies that were used including anti-HA tag (Abcam, ab9110) for HA-tagged Ikaros, anti-HDAC1 (Abcam, ab7028) and anti-histone modification antibodies: H3K4me3 (Abcam, ab8580), anti-H3K4me1(Abcam, ab8895), anti-H3K27ac (Abcam, ab4729) and anti-H3K27me3 (Millipore, 07-449). ChIP-seq libraries were created using ChIP-seq DNA sample prep kit (Illumina), size-selected and the 200-400bp fraction was extracted and purified. Libraries were sequenced at the High Throughput Genomics Center of University of Washington, Seattle and at Genome Sciences and bioinformatics core of Penn State University, Hershey, College of Medicine.

### CUT&Tag

CUT&Tag for W8 and Molt4 cells was performed according to EpiCypher CUTANA Direct-to-PCR CUT&Tag Protocol. Briefly, approximately 100k cells for each sample were collected. Nuclei were extracted by NE buffer, and were then immobilized to Concanavalin A conjugated paramagnetic beads (EpiCypher, 21-1401). Then the immobilized nuclei were incubated with primary then secondary antibodies. pAG-Tn5 (EpiCypher, 15-1017) were added to recognize antibodies which were the same for ChIP-seq, cleave target-DNA complex, and ligate sequencing adapters. Then DNA fragments were amplified by PCR, and purified by Agencourt AMPure XP beads (Beckman Coulter Genomics, #A63881). Sample libraries were sequenced on NovaSeq 6000 PE150 at Novogene, Sacramento, CA.

## Bioinformatics Analysis

### ChIP-seq Analysis

Both histone and transcription factor ChIP-seq data were analyzed using ENCODE3 pipeline. Briefly, Bowtie2 (ref. 1) (version 2.2.6) with default parameters to map fastq data to the mm9 reference genome. Samtools (ref. 2) (version 1.2) with MAPQ > 30 as a cutoff and PICARD (version 2.0.1) were used for further filtering. MACS2 (ref. 3) (version 2.1.1) was used to call peaks, followed by Benjamini-Hochberg procedure with q values of 0.05 as a cutoff for significance. For Transcription factor ChIP-seq, IDRR (ref. 4) (version 2.0.4) was further used to control the reproducibility between replicates with a cutoff value of 0.05. Finally, Narrowpeak files and p value bigwig signal files were used for downstream analysis.

Histone ChIP-seq peaks were used to define DNA elements. Promoters were defined as either H3K4me3 peak regions or 1.5kb upstream and downstream of TSS of genes defined in the annotation GTF file with GENCODE release version vM1. Poised Enhancers were defined as H3K4me1 peak regions that do not overlap with Promoter regions. Active Enhancers were defined as regions that have both H3K4me1 and H3K27Ac peaks. Gene Body regions were defined as annotation with the GENCODE release version vM1 but do not overlap with either Promoters or Enhancers. Ikaros regulated DNA elements were defined by overlapping Ikaros peaks with each of above-mentioned DNA elements. The remaining Ikaros binding regions were defined as Gene Desert regions.

De novo enhancers were defined as regions that gained H3K4me1 signals with Ikaros treatment as compared to wild type regardless of Ikaros binding or H3K27Ac signals. De novo activated enhancers were defined as a subset of de novo enhancers that also gained H3K27Ac signals. Ikaros regulated de novo enhancers are the subset of de novo enhancers that gained Ikaros binding when Ikaros-treated cells were compared with untreated cells.

To compare the functions of DNA elements across species, we used liftOver to convert the coordinates of the identified DNA elements from mouse genome (mm9) to human genome (hg38).

### Super-enhancers, Enhancers, LOCKs, and BGRDs

Super-enhancer identification was defined using HOMER (version 4.8). Firstly, peaks for H3K27Ac were found as described above. Then, peaks within 10 000 bp were combined together into larger regions. The H3K27Ac signals of these merged peak regions were then determined by the total input normalized number of reads as well as highest score and the total number of enhancer regions. Finally, intensity was potted against the rank of these enhancers, super-enhancers are identified as regions past the point where the tangent of slope is greater than 1, the rest would be typical enhancers.

Large organized chromatin lysine (K) domains (LOCKs) and broad genic repression domains (BGRDs) were identified following the original publications. To identify LOCKs, we first used MACS2 to identify H3K27me3 peaks in DN3 cells. And then we used CREAM algorithm to identify the peak clusters as LOCK (ref. 5). To identify BGRDs, after peak calling for H3K27me3 peaks, we retrieved the peaks that overlapped with each gene. We then plotted the maximal height of H3K27me3 peaks against the total width of H3K27me3 peaks for each gene. We finally used an H3K27me3 width of 120kb as the cutoff to define BGRDs (ref. 6).

### ATAC-seq analysis

ATAC-seq data were analyzed using the ENCODE3 ATAC-seq pipeline. Briefly, raw fastq data were firstly trimmed using CUTADAPT (ref. 7) (version 1.9.1) and then mapped to the hg19 reference genome with Bowtie2 (ref. 1) (version 2.2.6) using default parameters (-k –X2000 –local –-mm –-threads –x). Post-alignment filtering was achieved using Samtools (ref. 2) (version 1.2) with MAPQ > 30 as a cutoff, and removing duplicate reads using PICARD (version 2.0.1). MACS2 (ref. 3) (version 2.1.1) was used for calling peaks with q values of 0.05 as a cutoff. Finally, narrowpeak files and p value bigwig signal files were used for downstream analysis. Ikaros-regulated de novo open chromatin regions were defined by a gained ATAC-seq signal after treatment, as compared with untreated. Functional categories of these de novo open regions were determined by overlapping them with each category of DNA elements defined above.

### RNA-seq analysis

RNA-seq read data were aligned to mm9 reference genome using STAR (version 2.7.7) with –r 100 –no-coverage-search. FeatureCounts (ref. 8) (version 1.4.6) was then used to get read counts for each transcript. Symbol conversion was conducted through biomaRt (ref. 9) package (version 2.30.0). Data from three technical replicates were merged for subsequent analysis. Genes with less than 100 reads were filtered. The top-ranked differential regulated genes were determined by fold change. Welch’s two-sample t-test was used to determine the statistical significance between genes from two conditions. Volcano plots were generated in R using –log p-value plot against log fold change value. Significantly differentially expressed genes (p < 0.05) are color-coded: genes up-regulated by >2 fold are in orange, genes down-regulated by >2 fold are in purple, genes up or down regulated by <2 fold but with p < 0.05 are in red.

### Enhancer-target gene prediction

We used a recent developed algorithm Integrated Method for Predicting Enhancer Targets (IM-PET) to predict enhancer targets (ref. 10). IM-PET predicts enhancer-promoter by integrating four features using a Random Forest classifier. Features are derived from transcriptomic, epigenomic and genome sequence data, including enhancer and promoter activity correlation, TF and promoter activity correlation, enhancer and promoter sequence co-evolution and enhancer-to-promoter distance. We showed that IM-PET achieved significant improvement over other state-of-the-art methods. Further, based on our validation experiment using 3C-qPCR we showed that IM-PET has a comparable accuracy to that of the experimental 5C technology. Here, the input data for IM-PET included the genomic positions of predicted enhancers, RNA-Seq data and H3K4me1, H3K4me3, and H3K27Ac ChIP-Seq data for DN3 cells and Molt4 cells. Enhancer targets were predicted using a false discovery rate cutoff of 0.01.

### Heatmaps, boxplots, volcano plots and radar plots

Enrichment of transcription and Histone ChIP-seq signals in specific regions were analyzed and visualized using deeptools (ref. 11) (version 2.3.5) python package. Briefly, input normalized and p value filtered bigwig files generated from the ENCODE pipeline were used to compute a matrix for genomic regions of interest using computeMatrix function. In the matrix, each row is a genomic region and each column contains normalized chip-seq reads for each transcription factor/histone at a specific time point. The matrix was used as input for plotHeatmap function to generate heatmaps. Track view of examples for enhancers and super enhancers were generated using the UCSC genome browser with bigwig file as input. Volcano plots were generated in R using –log p-value against log fold change value. Significantly differential expressed genes are color coded as described in figure legends. Boxplots and radar plots were also generated in R.

### Motif enrichment analysis

Motif enrichment analysis was conducted using MEME Suite (ref. 12) (version 5.0.1) http://meme-suite.org<u>/</u> with database set to HOMOCOMO human (v11 core). Briefly, genomic sequence in fasta format for ChIP-seq peak regions were extracted using bedtools (version 2.25.0) from mm9 reference genome. MEME-ChIP function was then used to test the enrichment of known transcription factor-binding motifs in the database between target DNA sequences and randomly generated sequences. The length of each motif was set to a range from 3 to 10 bp. The minimum searching threshold was set as an E value of 0.05.

### GO term and pathway enrichment analysis

GO term and pathway enrichment analysis was achieved using The Database for Annotation, Visualization and Integrated Discovery (DAVID) (ref. 13) (version 6.8) https://david.ncifcrf.gov/. FDR cutoff 0.05 was used to define significant enrichment. For cis-regulatory regions, both Genomic Regions Enrichment of Annotations Tool (GREAT) (ref. 14) (version 3.0.0) http://great.stanford.edu/public/html/ and assignment with the closest TSS followed by a similar procedure as for gene approaches was used. Redar plot was conducted using –log10 of adjusted p values.

### GSEA analysis

Gene Set Enrichment Analysis (GSEA) software was downloaded from the Broad Institute website http://software.broadinstitute.org/gsea/index.jsp. Molecular Signatures Database (v6.1 MSigDB) was used to test enrichment for target genes with specific function. Briefly, the target set of genes were firstly ordered based on mean differential expression values of the two classes divided by the sum of the standard deviations. Then, an enrichment measure based on normalized Kolmogorov-Smirnov statistic was calculated for each gene set. The genes from a set with previously defined known function were computed by a running sum across all target genes. The ES enrichment score is defined as the maximum observed positive deviation of the running sum. ES is measured for every functional gene set considered. In this manuscript, after identifying Ikaros regulated genes, we tested it (a single gene set) for association with functions associated with T cells using GSEA.

### Data access

RNA-seq, ChIP-seq and ATAC-seq libraries were sequenced by Genome Sciences and bioinformatics core at Penn State University, Hershey, College of Medicine. All sequencing data are available as fastq-format files from the GEO archive under accession numbers GSE261180 and GSE261181

## Supplemental Figures

**Supplemental Figure 1.**
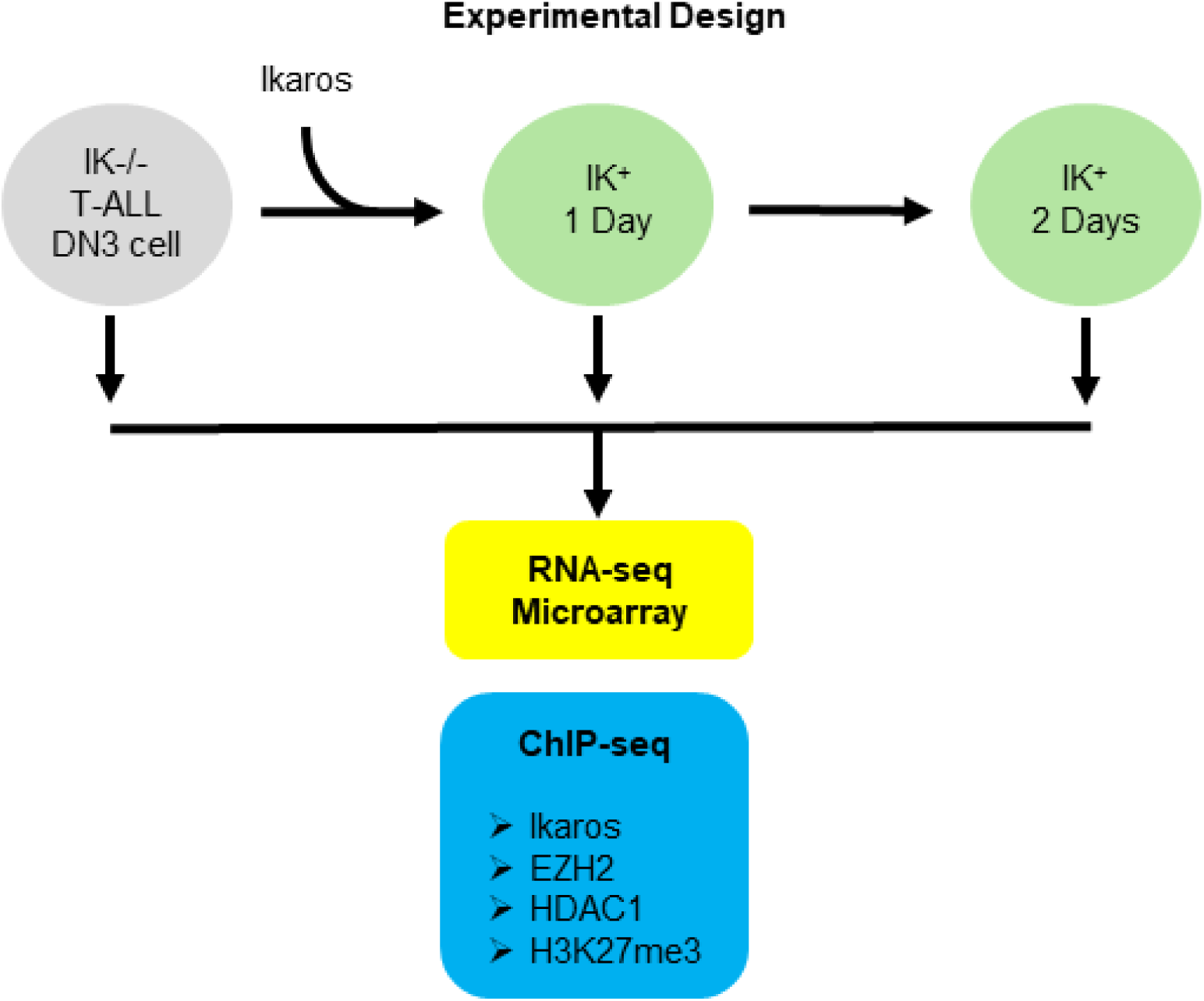
Experimental design. Ikaros (IK)-null murine T­ALL cells were transduced by Ikaros-containing retrovirus to express Ikaros. Cells were cultured for 2 Days following retroviral transduction. Cells were harvested prior to transduction (Day 0) and at daily time points over the 2 Day period (Day 1 and Day 2) for the indicated analyses.

**Supplemental Figure 2.**
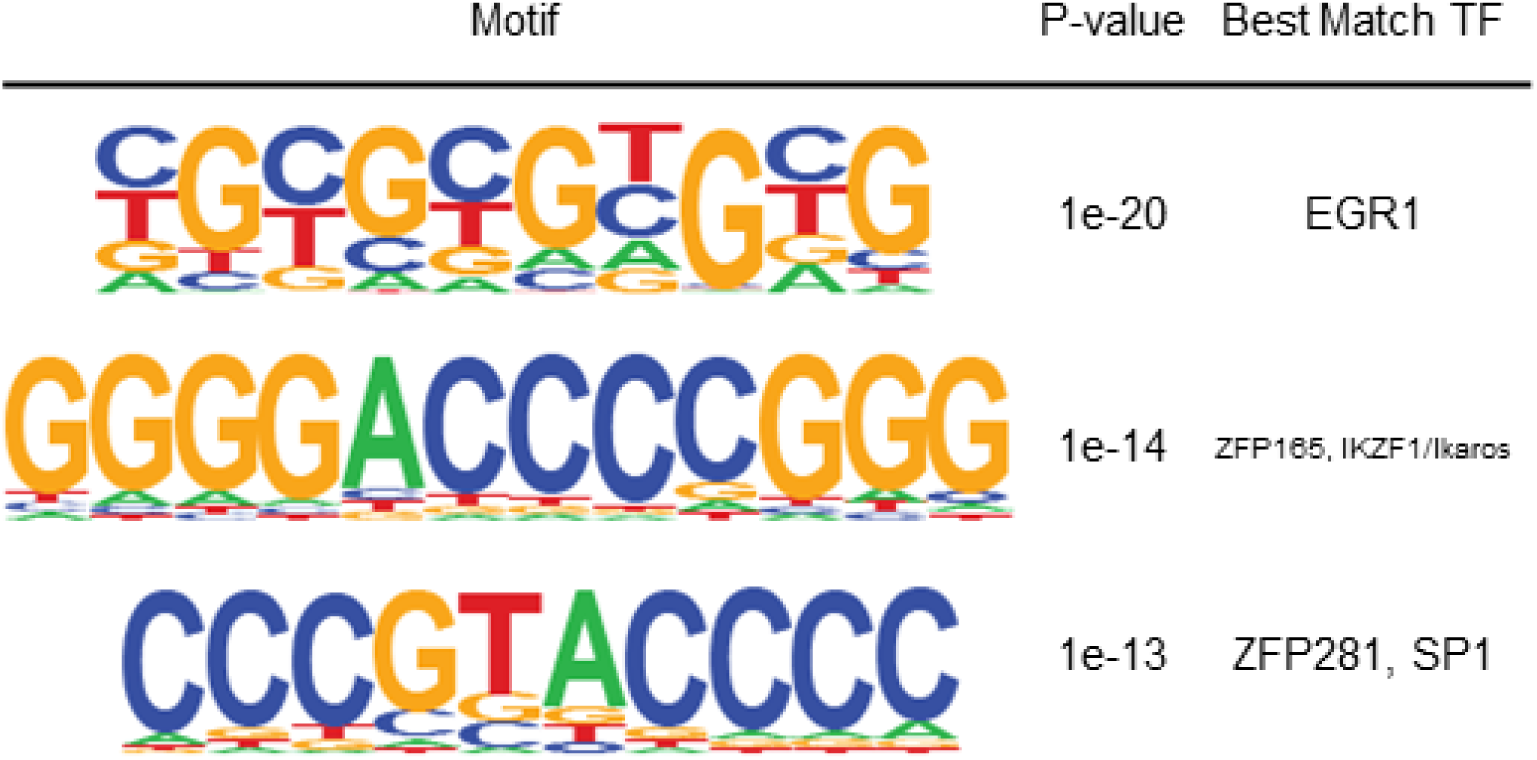
Motif enrichment analysis for de novo formed facultative heterochromatin (H3K27me3) peaks following Ikaros introduction at Day 1 MEME-ChIP was used to extract the enriched known transcription factor binding motifs from a large set of DNA sequences identified by ChlP-seq by searching against the JASPAR CORE database. Motif logo, significance of the motif, and names of transcription factors that bind the motifs are shown.

**Supplemental Figure 3.**
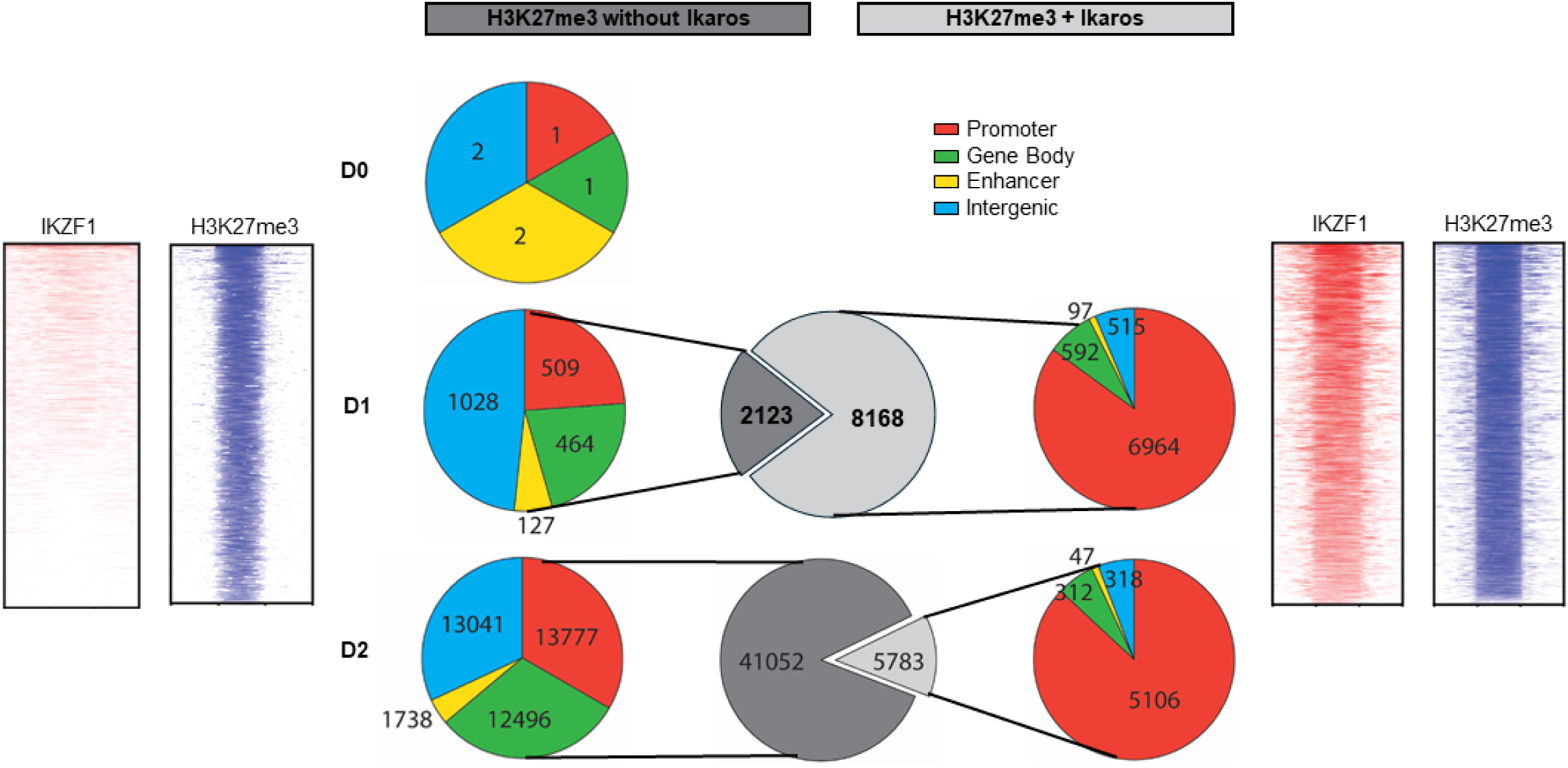
Dynamic changes in genome-wide distribution of Ikaros-induced facultative heterochromatin chromatin (H3K27me3). Dynamic changes in Ikaros binding-associated facultative heterochromatin (right-panel) and facultative heterochromatin formed following Ikaros re-introduction but not associated with Ikaros occupancy in untreated Ikaros-null T-ALL (Day 0) and for 2 Days following Ikaros re-introduction (Day 1 and Day 2). At Day 0 there are very few distinct H3K27me3 peaks. At Day 1 after transduction with Ikaros, the majority (8,168) of the de novo formed facultative heterochromatin (H3K27me3 peaks) are associated with Ikaros occupancy (right panel) as opposed to 2,128 H3K27me3 peaks which are not associated with Ikaros occupancy (left panel). In contrast, at Day 2 after transduction with Ikaros, there is a large increase in the total number of H3K27me3 peaks, with the majority of the new H3K27me3 peaks (41,052) not being associated with Ikaros occupancy (left lower panel), while 5,783 H3K27me3 peaks were associated with Ikaros binding (right lower panel). During both Days after Ikaros transduction, Ikaros-binding associated H3K27me3 are predominantly located at gene promoters, while H3K27me3 peaks not associated with Ikaros binding are relatively evenly distributed among promoters, gene body and intergenic regions, with a significant number of H3K27me3 peaks detected within enhancer regions in Day 2 (lower left panel – yellow). Heat maps for Ikaros binding-associated (right) and Ikaros binding-independent (left) H3K27me3 peaks are shown.

**Supplemental Figure 4.**
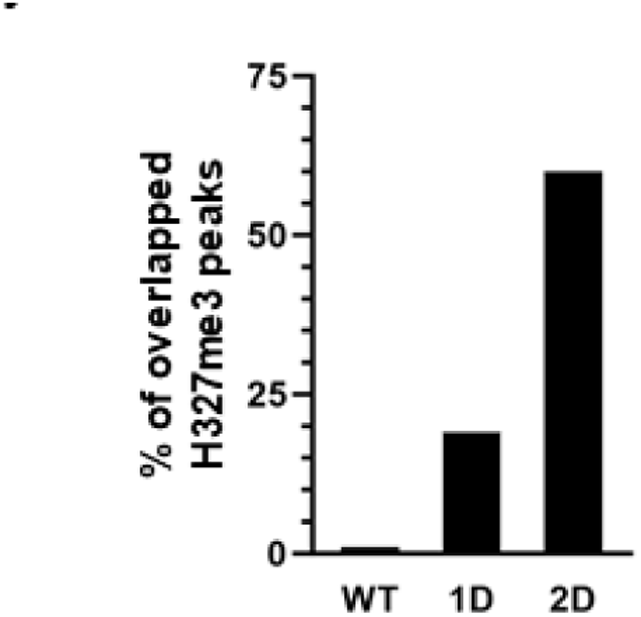
Re-introduction of Ikaros into Ikaros-null T­ALL progressively restores normal H3K27me3 landscape. Results show that Ikaros re-introduction over 2 Days period significantly restores H3K27me3 landscape, which at the end of 2 Days overlaps with 60% of H3K27me3 peaks in normal thymocytes at DN3 stage of differentiation.

**Supplemental Figure 5.**
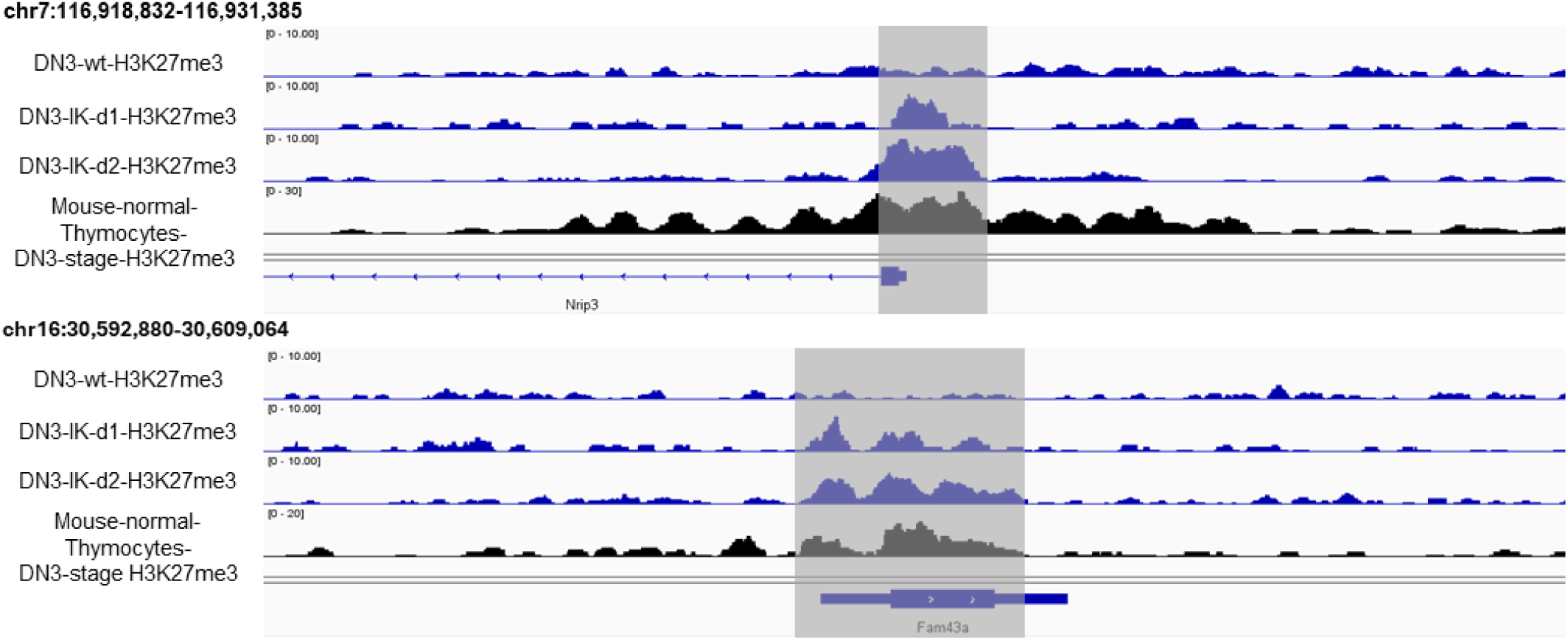
Example of the dynamic restoration of normal H3K27me3 landscape in IK-null T-ALL following Ikaros re-introduction. Comparison of published H3K27me3 landscape in normal thymocytes at DN3 stage of differentiation (bottom panels) with IK-null T-ALL and following Ikaros re­introduction in Day 1 and 2 (top 3 panels) – example. Results show that normal H3K27me3 landscape is erased in IK-null T-ALL, but Ikaros re-introduction restores normal (physiological) H3K27me3 landscape.

**Supplemental Figure 6.**
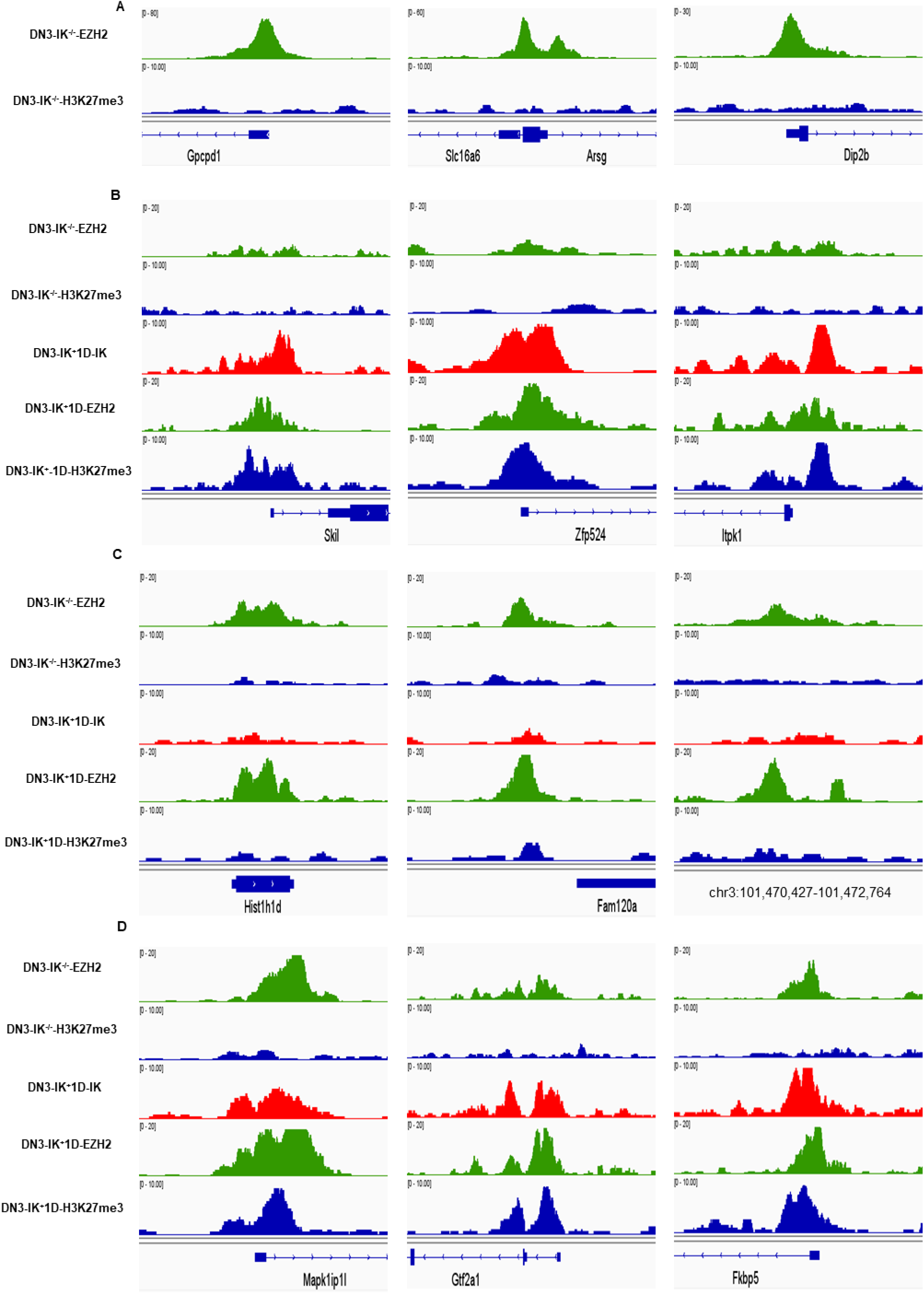
(A) Examples of EZH2 DNA binding in Ikaros-null T-ALL without H3K27me3 occupancy; (B) Recruitment of EZH2 by Ikaros results in de novo formation of EZH2-lkaros DNA-binding complexes in IK*^D^ay 1 following Ikaros re-introduction, resulting in formation of H3K27me3 (Day 0 EZH2 not binding, no H3K27me3; IK*Day 1 Ikaros binding, EZH2 binding H3K27me3 formation); (C) EZH2 binding to DNA without concomitant Ikaros binding, 1 Day following Ikaros re-introduction is not associated with H3K27me3 (Day 1 EZH2 peak, no Ikaros peak, no H3K27me3 peak). (D) Sites occupied solely by EZH2 in Ik-null T-ALL, and not being enriched in H3K27me3, following formation of lkaros-EZH2 complexes in Day #1, result in H3K27me3 enrichment – EZH2 binding in Day 0; no H3K27me3 Day 0, Ikaros binding in Day 1, EZH2 binding in Day 1 and H3K27me3 peak in Day 1;

**Supplemental Figure 7.**
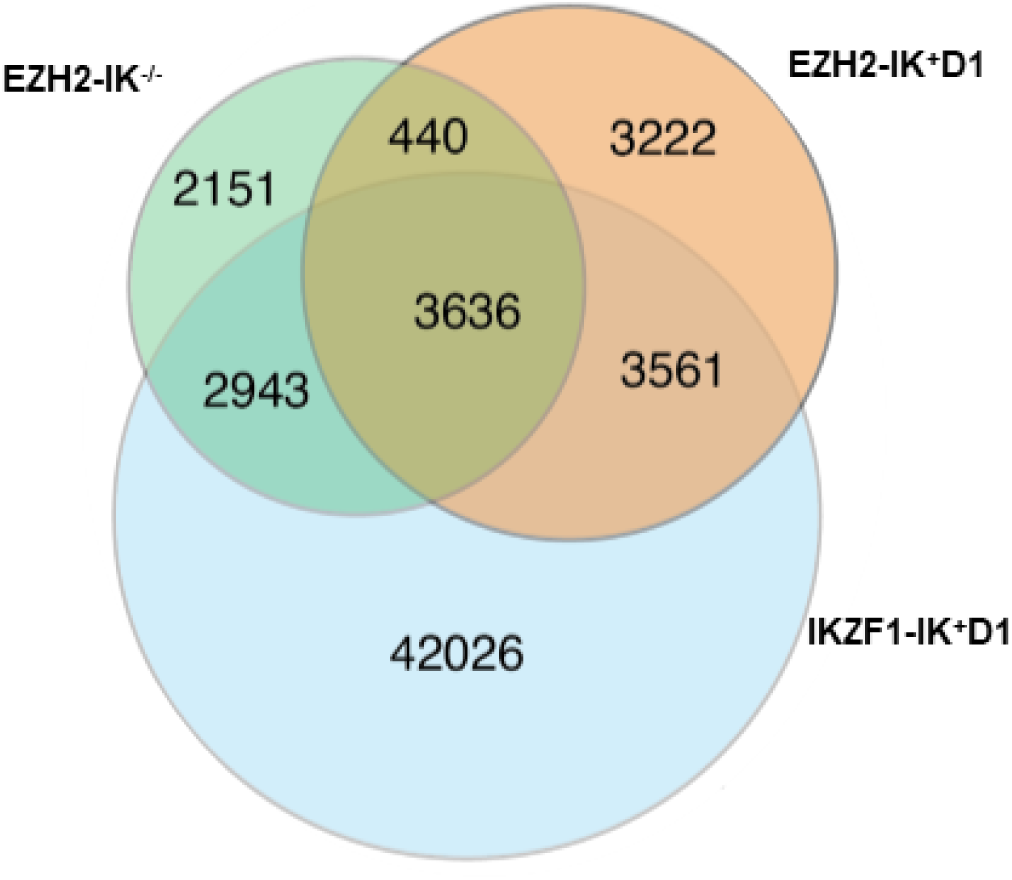
Genome distribution of EZH2 ChlP-seq peaks in Ikaros-null T-ALL (Day 0) and 1 Day after Ikaros transduction (D1), as well as Ikaros ChlP-seq peaks 1 Day after Ikaros transduction. Results show that Ikaros binding results in redistribution of EZH2 genome occupancy either by direct recruitment by Ikaros (3,561 peaks), or via Ikaros-independent manner (3,222 peaks). A large number (3,636) EZH2 binding sites, that are occupied in Ikaros-null T-ALL are bound by Ikaros following Ikaros transduction, resulting in formation of EZH2-lkaros complexes.

**Supplemental Figure 8.**
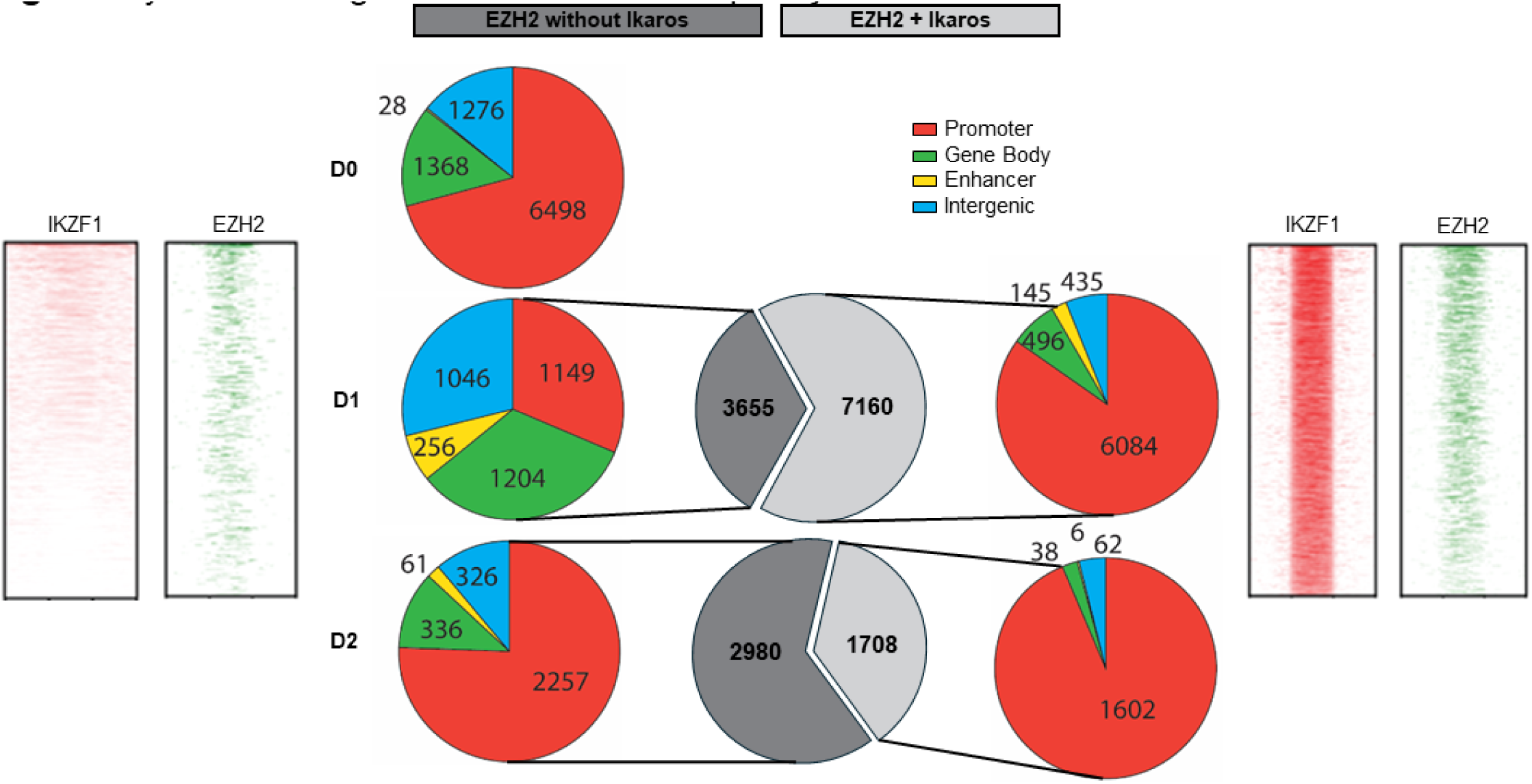
Dynamic changes EZH2-only and EZH2-lkaros occupied sites classified by function of the DNA element Dynamic changes of EZH2 DNA binding sites in Ikaros-null T-ALL (top panel) and EZH2-lkaros and facultative heterochromatin formed following Ikaros re-introduction but not associated with Ikaros occupancy in untreated Ikaros-null T-ALL (Day 0) and for 2 s following Ikaros re-introduction (day 1 and day 2). At Day 0 there are very few distinct H3K27me3 peaks. At Day 1 after transduction with Ikaros, the majority (Day8,168) of the de novo formed facultative heterochromatin (H3K27me3 peaks) are associated with Ikaros occupancy (right panel) as opposed to 2,128 H3K27me3 peaks which are not associated with Ikaros occupancy (left panel). In contrast, at Day 2 after transduction with Ikaros, there is a large increase in the total number of H3K27me3 peaks, with the majority of the new H3K27me3 peaks (41,052) not being associated with Ikaros occupancy (left lower panel), while 5,783 H3K27me3 peaks were associated with Ikaros binding (right lower panel). During both Days after Ikaros transduction, Ikaros-binding associated H3K27me3 are predominantly located at gene promoters, while H3K27me3 peaks not associated with Ikaros binding are relatively evenly distributed among promoters, gene body and intergenic regions, with a significant number of H3K27me3 peaks detected within enhancer regions in Day 2 (lower left panel – yellow). Heat maps for Ikaros binding-associated (right) and Ikaros binding-independent (left) H3K27me3 peaks are shown.

**Supplemental Figure 9.**
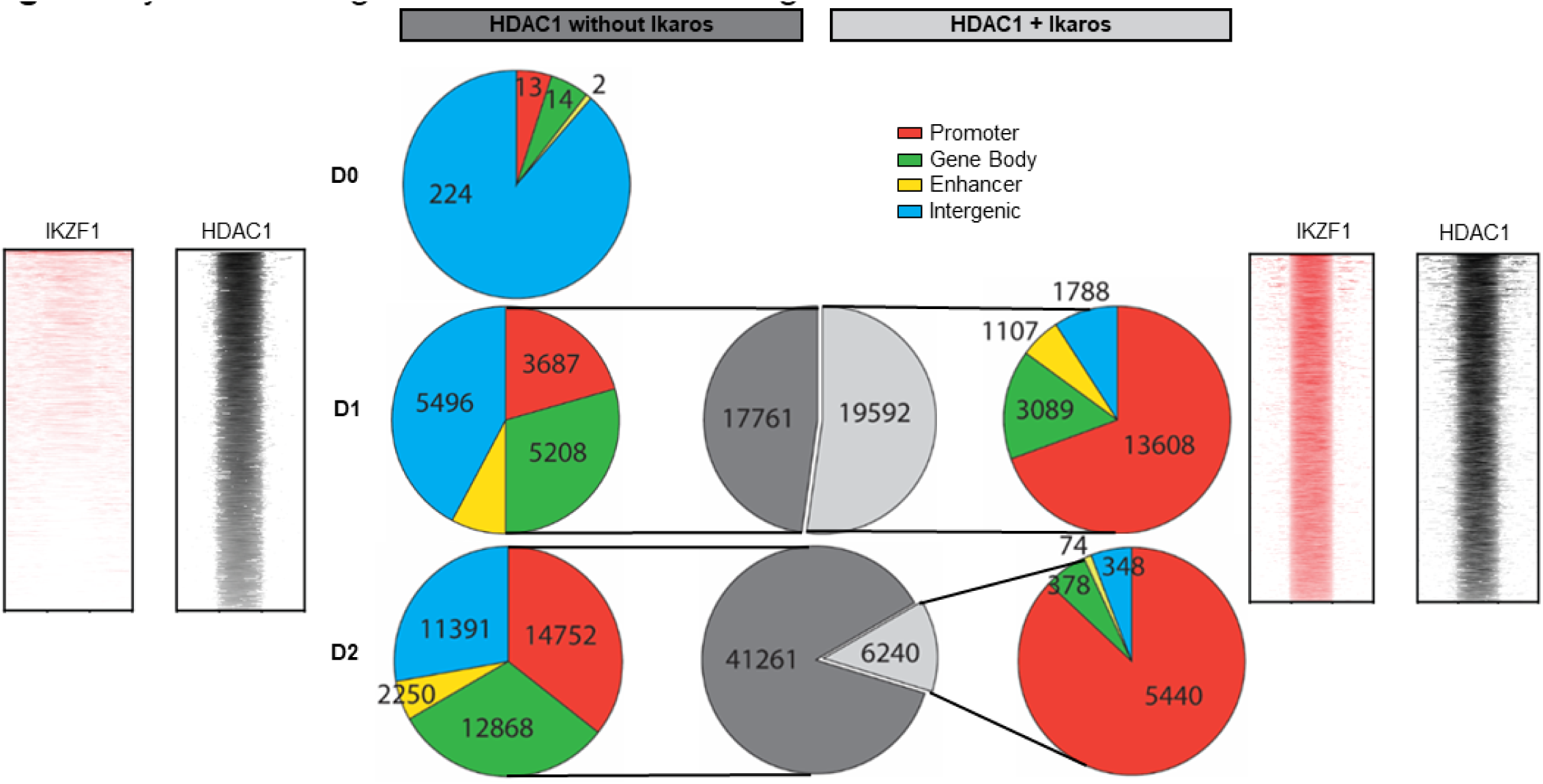
Dynamic changes HDAC1-only and HDAC1-lkaros occupied sites classified by function of the DNA element Dynamic changes in DNA occupancy of HDAC1 without Ikaros (left) and HDAC1-Ikaros complexes (right) are show in Ikaros-null T-ALL (Day 0) and for 2 Days following Ikaros re-introduction (Day 1 and Day 2). At Day 0 there are very few HDACIpeaks. At Day 1 after transduction with Ikaros, over half (19,592) of the HDAC1 peaks are associated with Ikaros occupancy (right panel) as opposed to 17,761 HDAC1 peaks which are not associated with Ikaros occupancy (left panel). In contrast, at Day 2 after transduction with Ikaros, there is a large increase in the total number of HDAC1 peaks, with the majority of the new HDAC1 peaks (41,261) not being associated with Ikaros occupancy (left lower panel), while 6,240 HDAC1 peaks were associated with Ikaros binding (right lower panel). During both Days after Ikaros transduction, lkaros-HDAC1 complexes are predominantly located at gene promoters, while HDAC1 peaks not associated with Ikaros binding are relatively evenly distributed among promoters, gene body and intergenic regions, with a significant number of HDAC1 peaks detected within enhancer regions in Day 2 (lower left panel – yellow). Heat maps for Ikaros binding-associated (right) and Ikaros binding-independent (left) HDAC1 peaks are shown.

**Supplemental Figure 10.**
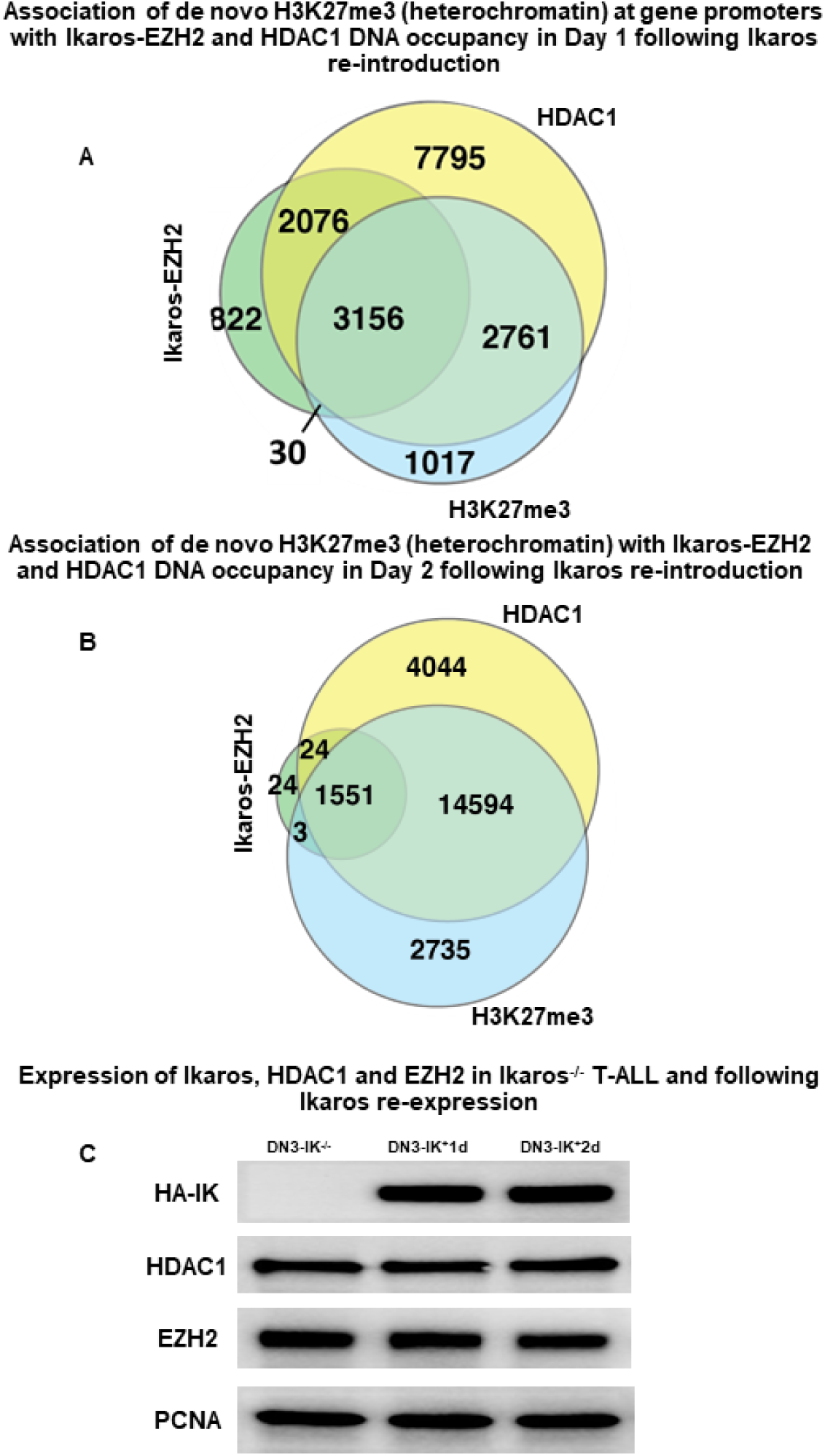
Association of Ikaros, EZH2 and HDAC1 with de novo formation of H3K27me3 at promoters. Results demonstrate a predominant role of HDAC1 in de novo formation of H3K27me3 (A) In Day 1 following Ikaros expression, formation of H3K27me3 at promoters is similarly associated with Ikaros, EZH2 and HDAC1 co-occupancy (3,156 peaks) and with HDAC1-only occupancy (2,761 peaks). (B) In contrast, in Day2 following Ikaros re-introduction, most of the H3K27me3 peaks at promoters (14,594) are associated with HDAC1-only DNA binding. Notably, no H3K27me3 peaks are associated with EZH2-only DNA binding, further supporting the hypothesis that EZH2 H3K27me3 methyltransferase activity requires Ikaros and/or HDAC1 DNA occupancy. (C) Western blot analysis shows that expression level of HDAC1 and EZH2 are similar in DN3 IK’– and following Ikaros re-expression. PCNA expression is used as a loading control.

**Supplemental Figure 11.**
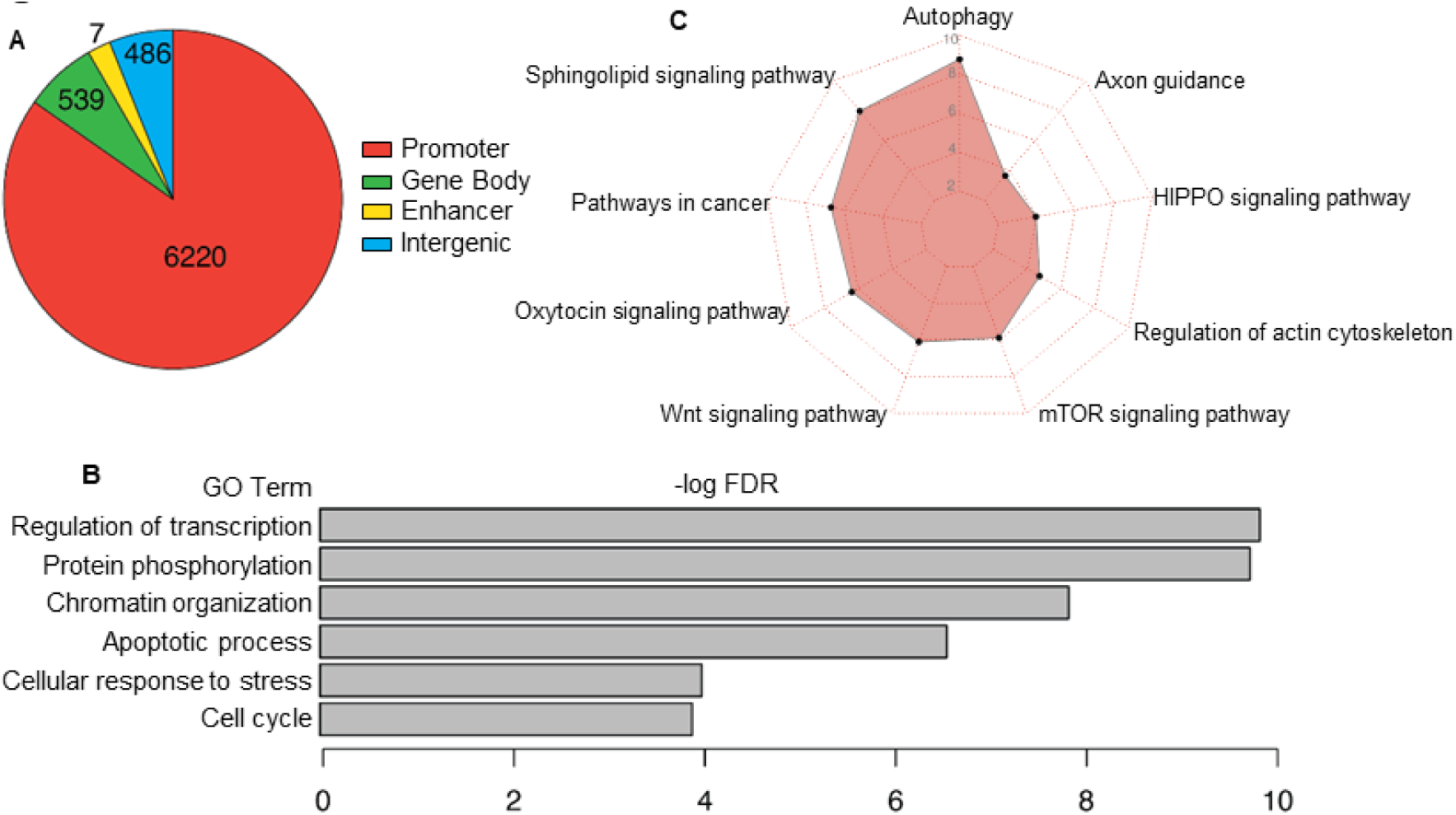
EZH2 target genes that are regulated by lkaros/HDAC1 induced epigenetic switch. (A) Distribution of de novo induced facultative heterochromatin (H3K27me3) at EZH2-occupied sites via Ikaros and HDAC1-induced epigenetic switch following Ikaros re-introduction to Ikaros-null T-ALL. (B-C) Gene ontology and pathway enrichment analysis of EZH2 target genes that are regulated by Ikaros and HDAC1-induced epigenetic switch. Many pathways that are regulated by lkaros/HDAC1-induced epigenetic switch are known for their oncogenic activity. Results suggest that Ikaros-null T-ALL results in functional inactivation of EZH2, lack of H3K27me3 occupancy, and activation of the above oncogenic pathways. Ikaros/HDAC1-induced switch to H3K27me3 occupancy results in repression of oncogenic pathways and tumor suppression.

**Supplemental Figure 12.**
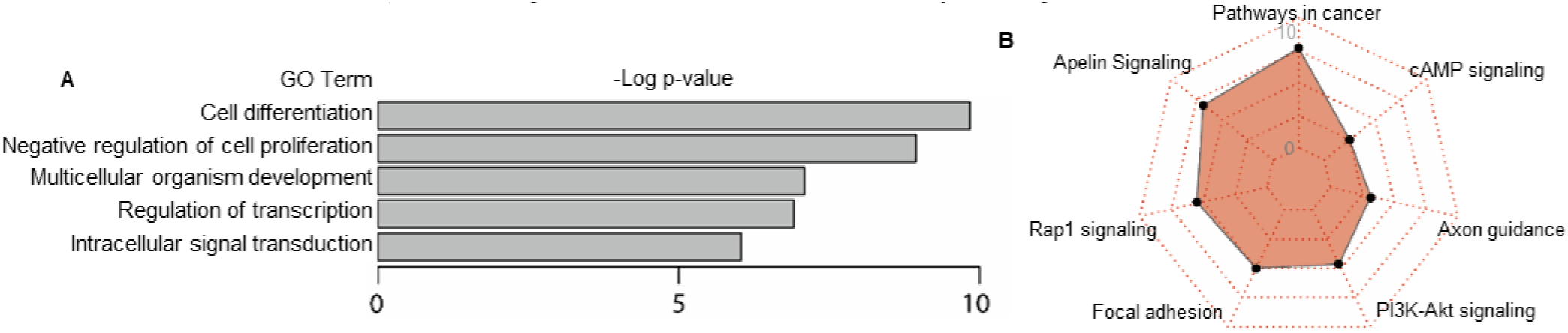
(A) Gene ontology and (8) pathway enrichment analysis of the genes that are regulated by active enhancers occupied by lkaros but not by HDAC1

**Supplemental Figure 13.**
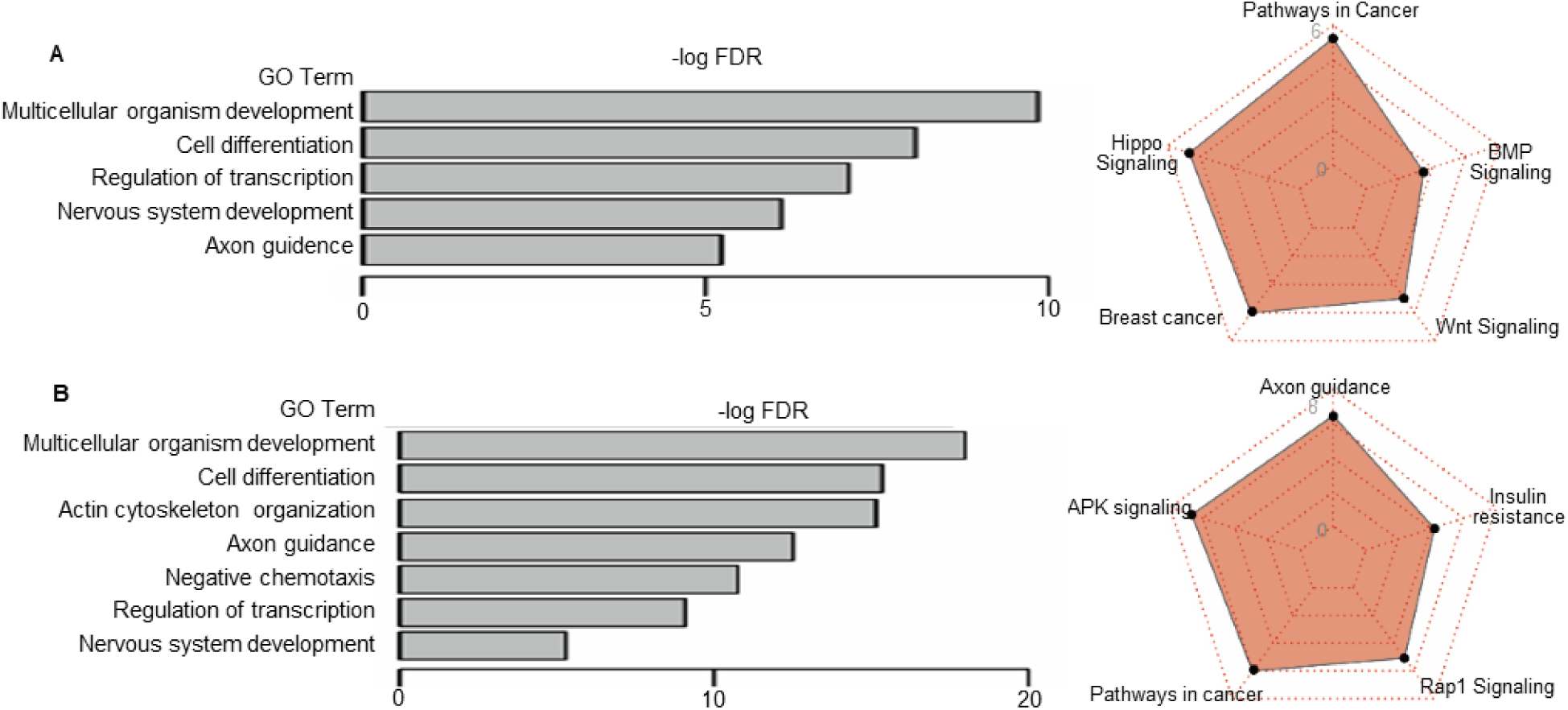
Gene ontology (left) and pathway enrichment analysis (right) of the genes that are regulated by active enhancers occupied by both HDAC1 but not lkaros in (A) Day 1 and (B) Day 2 following lkaros re-introduction

**Supplemental Figure 14.**
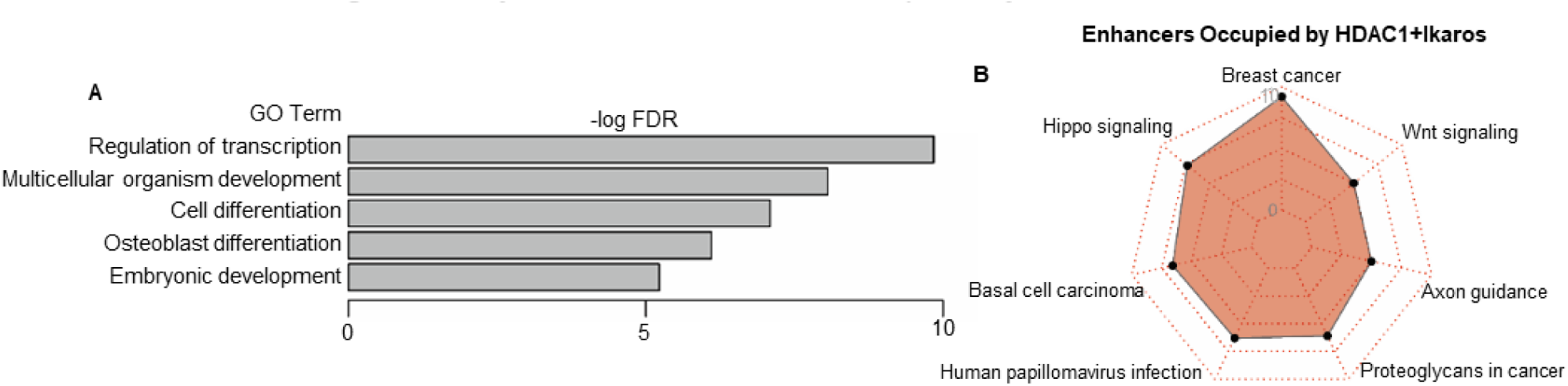
(Al. Gene ontology and (Bl pathway enrichment analysis of the genes that are regulated by active enhancers occupied by both lkaros and HDAC1

**Supplemental Figure 15.**
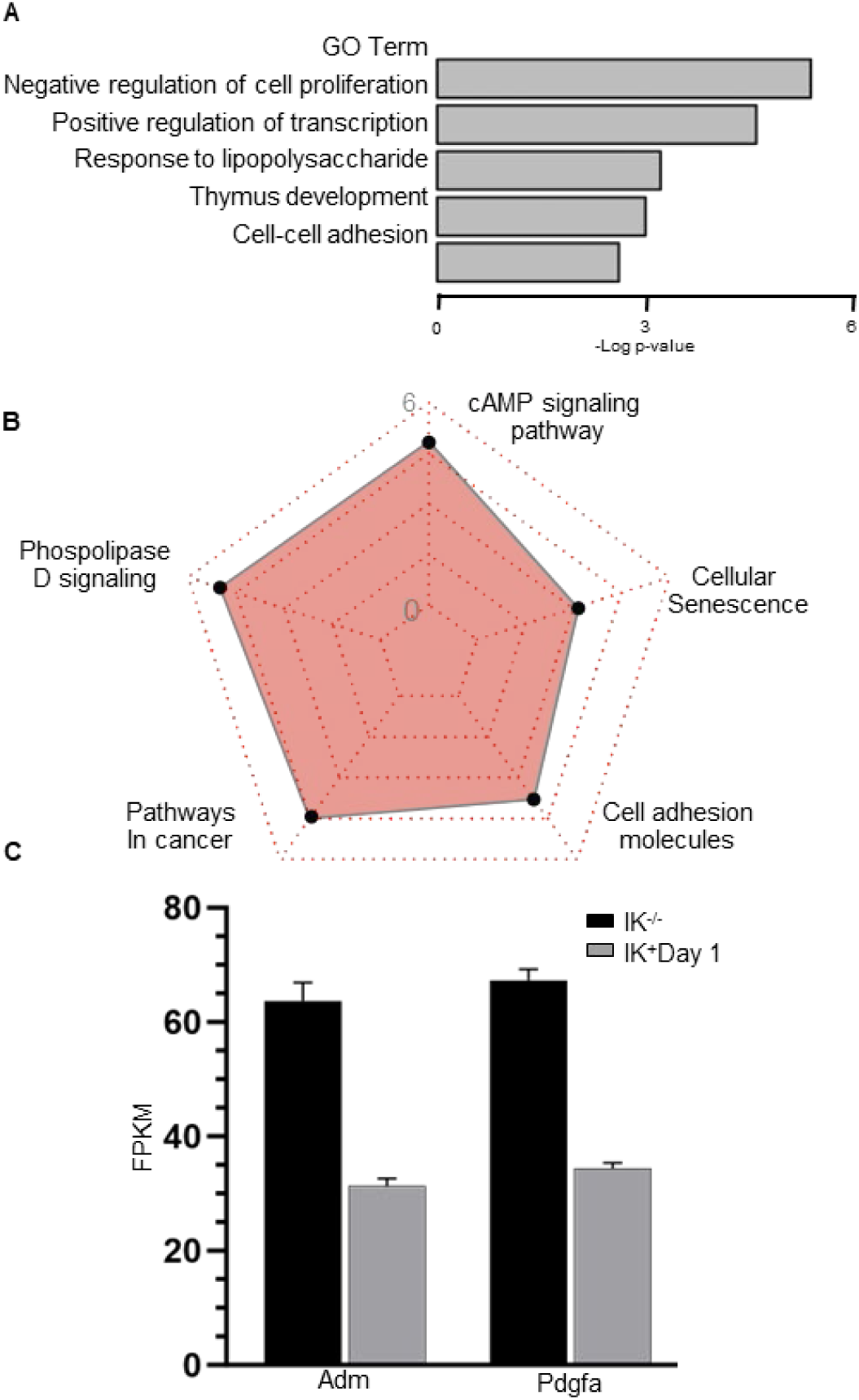
(A) Gene ontology (B) pathway enrichment analysis of genes regulated by Ikaros^7^-T-ALL, which became silenced by Ikaros and/or HDAC1 following Ikaros re-epression. (C) examples of gene repression due to enhancer silencing by Ikaros and HDAC1 following Ikaros re-expression.

**Supplemental Figure 16.**
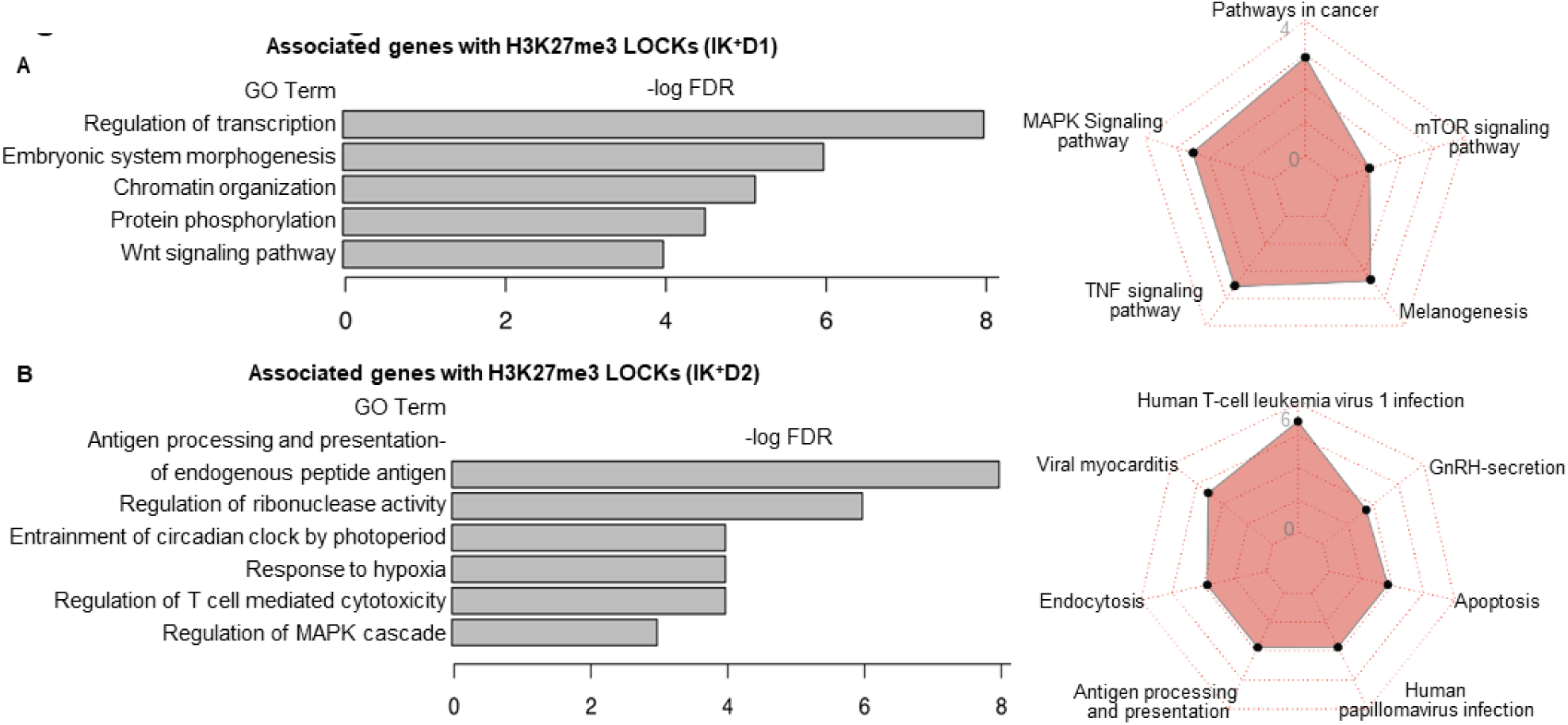
Gene ontology (left) and pathway enrichment analysis (right) of the genes that are found within the H3K27me3 LOCKs in (A) Day 1 and (B) Day 2 following lkaros re-expression

**Supplemental Figure 17.**
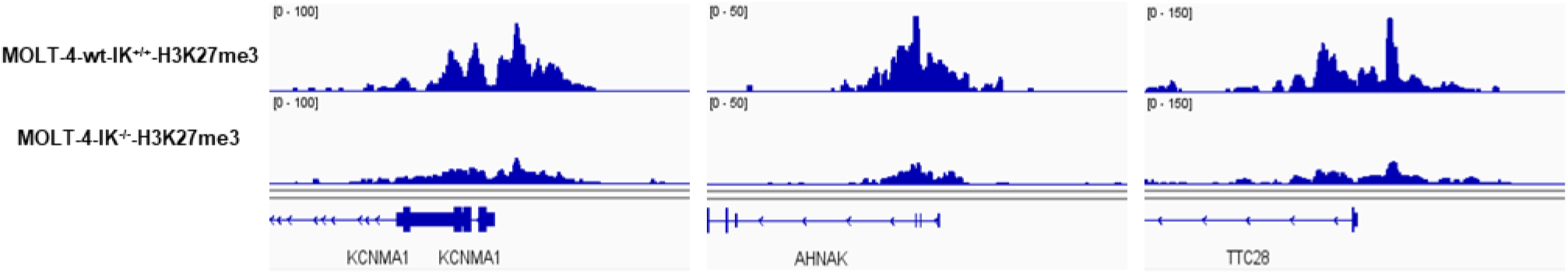
Example of the loss of H3K27me3 in human T-ALL associated with absence of Ikaros. Comparison of H3K27me3 CUT&Tag landscape in human T-ALL MOLT-4 cells, which express wild type Ikaros (top panel) with H3K27me3 distribution obtained by CUT&Tag on MOLT-4 cells with CRISPR-induced IKZF1 knockout. Results show that Ikaros knockout results in severely reduced H3K27me3 landscape in human T­ALL

**Supplemental Figure 18.**
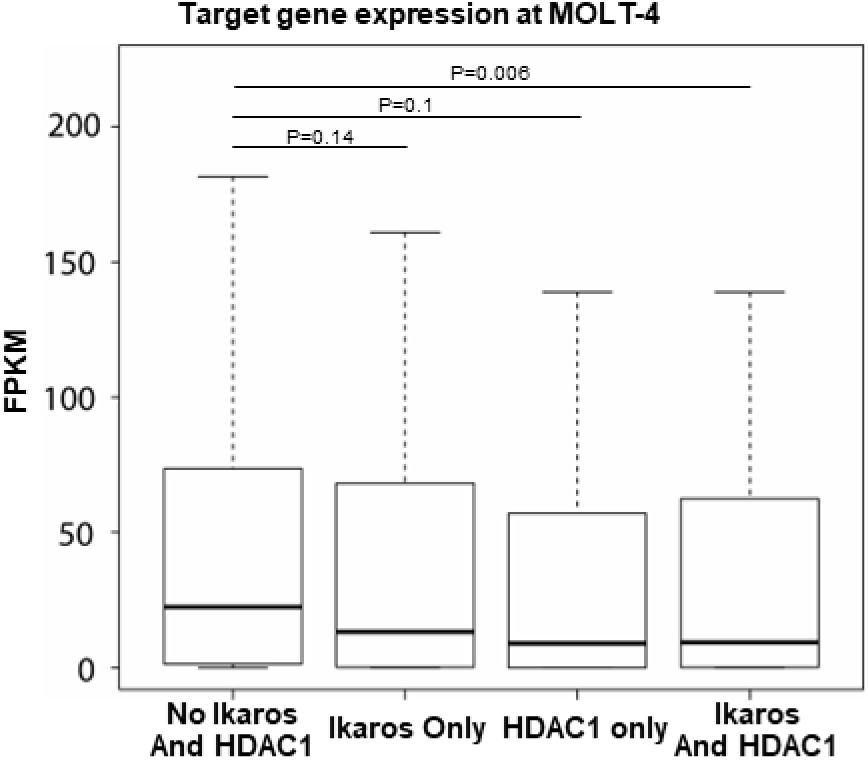
Ikaros and HDAC1 occupancy represses active enhancers in human T-ALL. Boxplot shows expression of genes regulated by the active enhancers in human T-ALL not bound by Ikaros and/or HDAC1 (left) and occupied by Ikaros and/or HDAC1. Boxplot: center line represents median, boxes show first and third quartiles, whiskers extend to the most extreme data points that are no more than 1.5-fold of the interquartile range from the box.

**Supplemental Figure 19.**
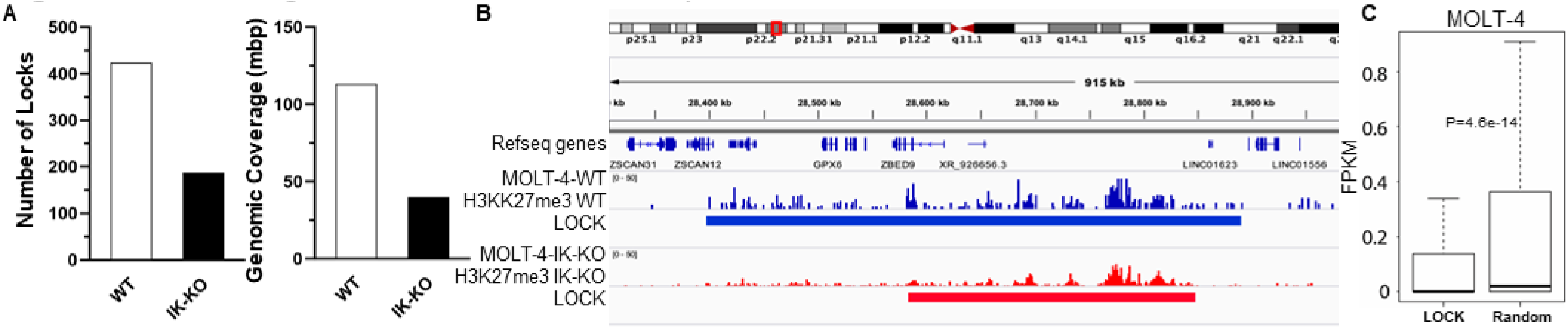
Ikaros regulates formation and expansion of H3K27me3 LOCKs in human T-ALL. (A) Number (left) and genomic coverage (right) of H3K27me3 LOCKs in human T-ALL with wildtype Ikaros (IK-wt) and Ikaros knockout (IK-KO) (B) Example of the genomic expansion of the H3K27me3 LOCKs in human T-ALL with wildtype Ikaros (IK-wt) vs. human T-ALL Ikaros knockout (IK-KO). (C) Negative regulation of gene by LOCKs. Expression of genes located within IOCK compared to the expression of the genes not regulated by LOCKS.

**Supplemental Figure 20.**
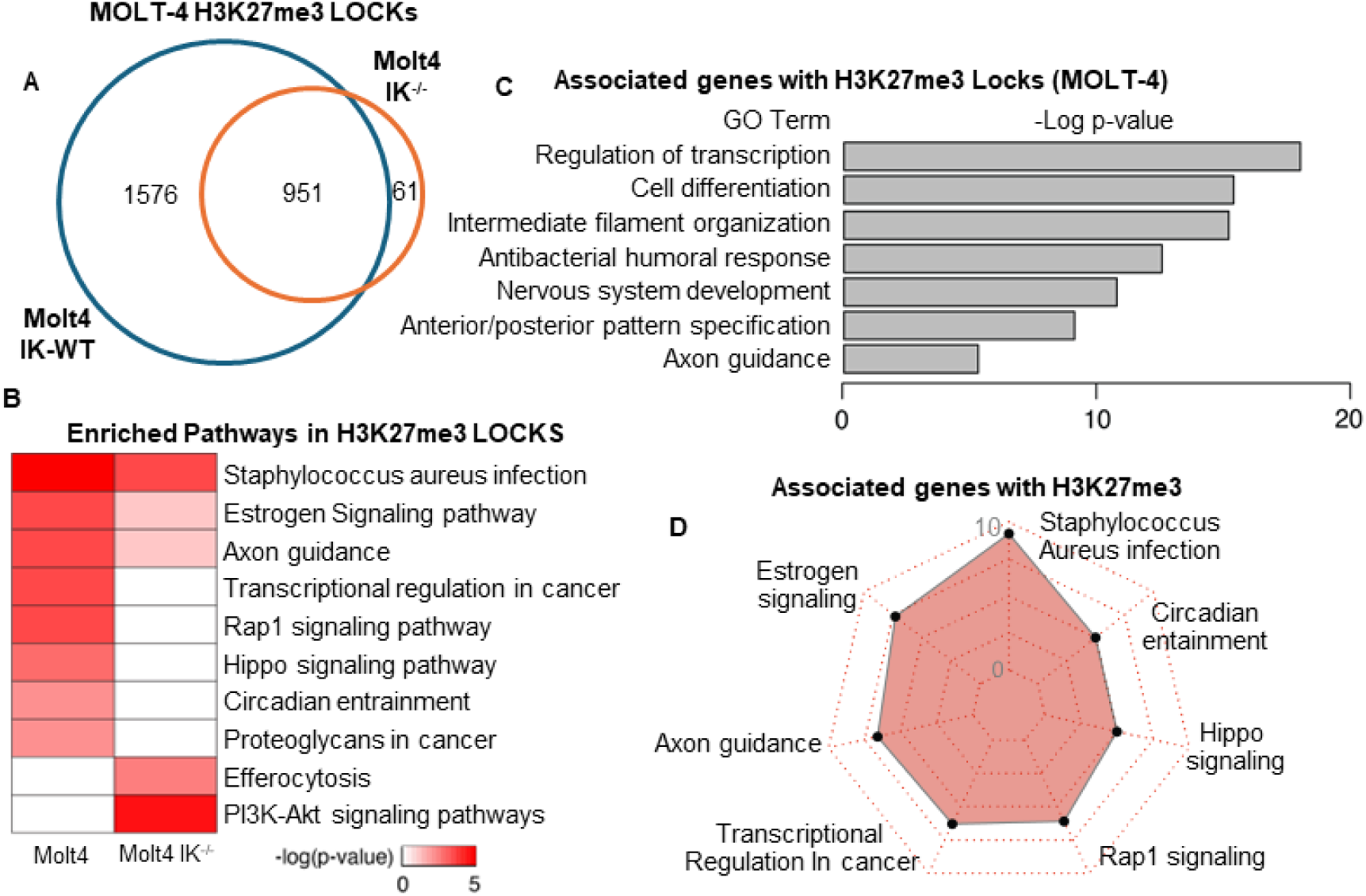
Ikaros regulates large set of genes in human T-ALL via formation of H3K27me3 LOCKs A) Number of genes regulated by the H3K27me3 LOCKs in human T-ALL with wildtype Ikaros (IK-wt) and Ikaros knockout (IK-KO). (B) Gene set enrichment analysis on the genes found within the H3K27me3 LOCKs in human T­ALL with wildtype Ikaros (IK-wt) and Ikaros knockout (IK-KO). (C) Gene ontology and (D) Pathway analysis of the genes found within H3K27me3 LOCKs in human T-ALL with wildtype Ikaros.

**Supplemental Figure 21.**
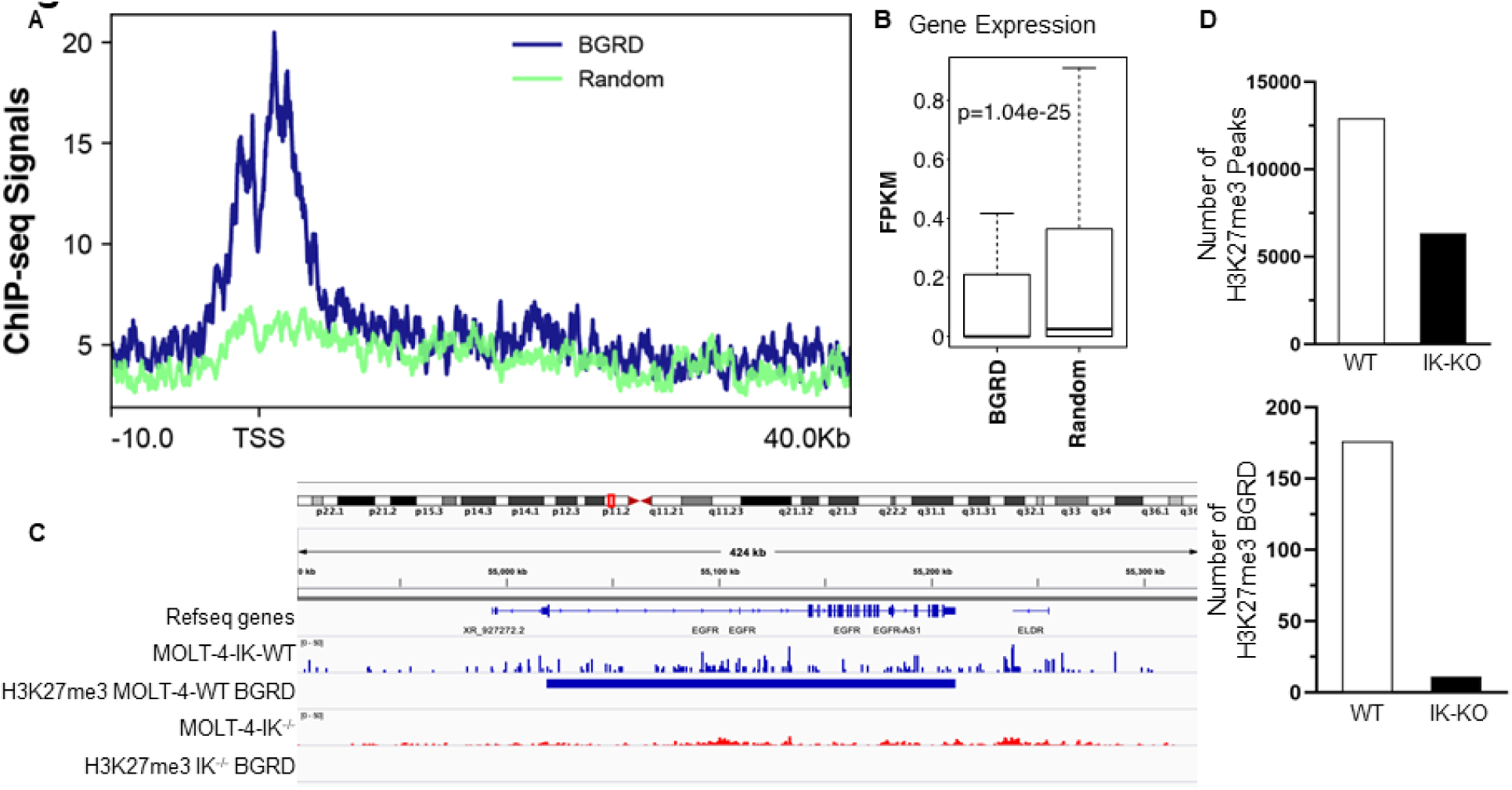
Ikaros regulate formation of H3K27me3 BGRDs in human T-ALL. (A) Average ChlP-seq signal value for H3K27me3 plotted around TSS of the genes within BGRD region vs. the random genes (B) Boxplot shows the expression value of the genes (by RNA-seq) located within BGRD vs. random genes. Center line is median, while boxes show first and third quartiles, with whiskers extending to the most extreme data points that are no more than 1.5-fold of the interquartile range from the box. (C) Example of BGRD which include EGFR oncogene in human T-ALL with wildtype Ikaros (IK-WT) vs. human T-ALL with Ikaros knockout (IK-KO). (D) List of the oncogenes and tumor suppressor genes associated with H3K27me3 BGRDs in human T-ALL with wildtype Ikaros (IK-WT) vs. human T-ALL with Ikaros knockout (IK-KO).

**Supplemental Figure 22.**
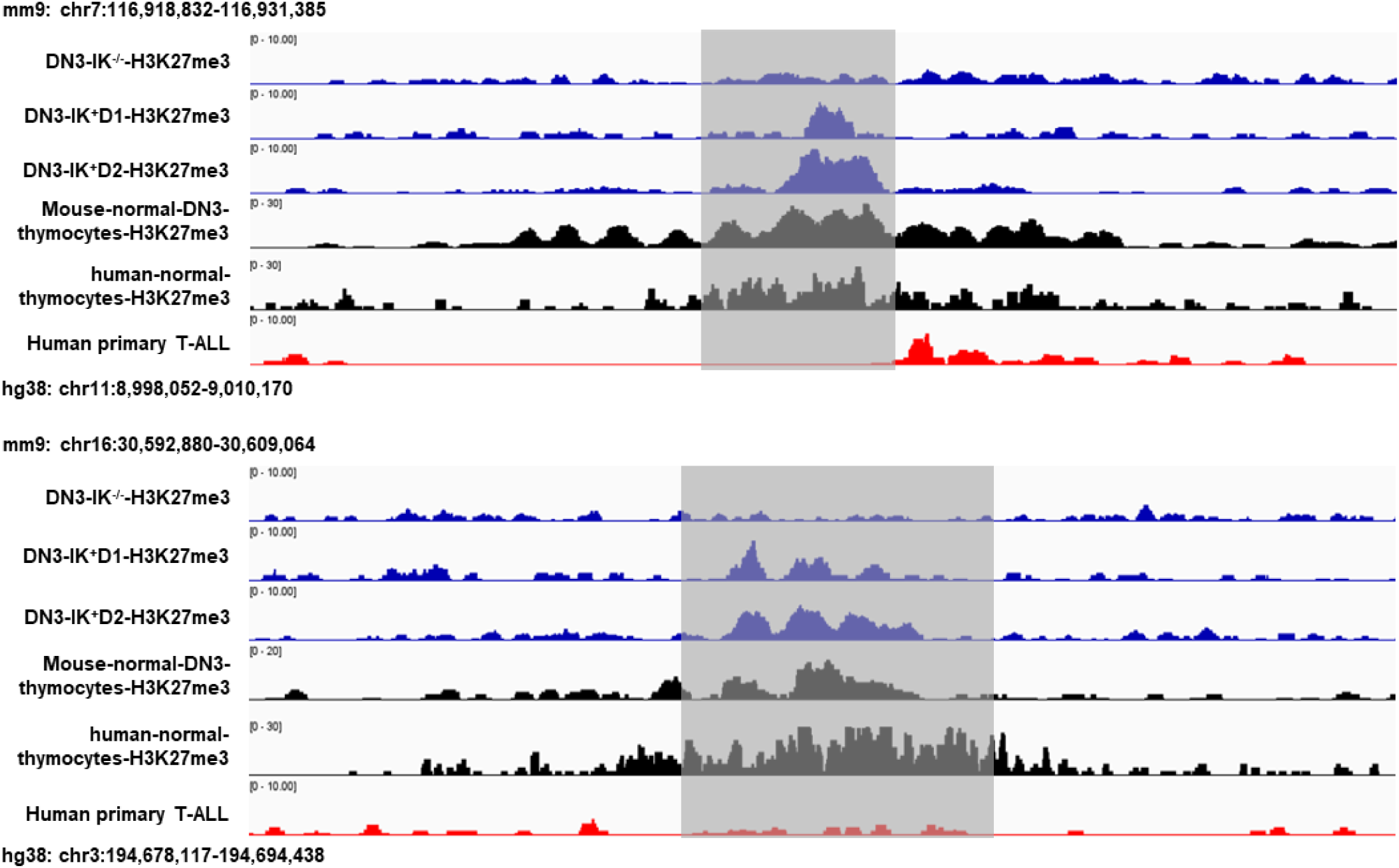
Example of conservation of H3K27me3 heterochromatin in mouse Ikaros-null and human T-ALL (DN3 and W8), normal mouse and human thymocytes, and the role of Ikaros in re-establishing normal H3K27me3 landscape. Results show loss of H3K27me3 signature in mouse and human T-ALL (top and bottom panel). Re-introduction of Ikaros into mouse Ikaros-null T-ALL results in partial re-establishment of the normal H3K27me3 landscape that is observed in mouse and human thymocytes.

## References

1. Georgopoulos K, Moore DD, Derfler B. Ikaros, an early lymphoid-specific transcription factor and a putative mediator for T cell commitment. Science. 1992;258(5083):808–12.

2. Lo K, Landau NR, Smale ST. LyF-1, a transcriptional regulator that interactswith a novel class of promoters for lymphocyte-specific genes. Mollecular Cellular Biology. 1991;11:5229–43.

3. Georgopoulos K, Bigby M, Wang JH, Molnar A, Wu P, Winandy S, et al. The Ikaros gene is required for the development of all lymphoid lineages. Cell. 1994;79(1):143–56.

4. Mullighan CG, Goorha S, Radtke I, Miller CB, Coustan-Smith E, Dalton JD, et al. Genome-wide analysis of genetic alterations in acute lymphoblastic leukaemia. Nature. 2007;446(7137):758–64.

5. Mullighan CG, Su X, Zhang J, Radtke I, Phillips LA, Miller CB, et al. Deletion of IKZF1 and prognosis in acute lymphoblastic leukemia. The New England journal of medicine. 2009;360(5):470–80.

6. Goldman FD, Gurel Z, Al-Zubeidi D, Fried AJ, Icardi M, Song C, et al. Congenital pancytopenia and absence of B lymphocytes in a neonate with a mutation in the Ikaros gene. Pediatric blood & cancer. 2012;58(4):591–7.

7. Kuiper RP, Schoenmakers EF, van Reijmersdal SV, Hehir-Kwa JY, van Kessel AG, van Leeuwen FN, et al. High-resolution genomic profiling of childhood ALL reveals novel recurrent genetic lesions affecting pathways involved in lymphocyte differentiation and cell cycle progression. Leukemia. 2007;21(6):1258–66.

8. Kuiper RP, Waanders E, van der Velden VH, van Reijmersdal SV, Venkatachalam R, Scheijen B, et al. IKZF1 deletions predict relapse in uniformly treated pediatric precursor B-ALL. Leukemia. 2010;24(7):1258–64.

9. van der Veer A, Waanders E, Pieters R, Willemse ME, Van Reijmersdal SV, Russell LJ, et al. Independent prognostic value of BCR-ABL1-like signature and IKZF1 deletion, but not high CRLF2 expression, in children with B-cell precursor ALL. Blood. 2013.

10. Kuehn HS, Nunes-Santos CJ, Rosenzweig SD. IKAROS-Associated Diseases in 2020: Genotypes, Phenotypes, and Outcomes in Primary Immune Deficiency/Inborn Errors of Immunity. J Clin Immunol. 2021;41(1):1–10.

11. Kuehn HS, Nunes-Santos CJ, Rosenzweig SD. Germline IKZF1 mutations and their impact on immunity: IKAROS-associated diseases and pathophysiology. Expert Rev Clin Immunol. 2021;17(4):407–16.

12. Raca G, Abdel-Azim H, Yue F, Broach J, Payne JL, Reeves ME, et al. Increased Incidence of IKZF1 deletions and IGH-CRLF2 translocations in B-ALL of Hispanic/Latino children-a novel health disparity. Leukemia. 2021.

13. Zhang J, Ding L, Holmfeldt L, Wu G, Heatley SL, Payne-Turner D, et al. The genetic basis of early T-cell precursor acute lymphoblastic leukaemia. Nature. 2012;481(7380):157–63.

14. Kim J, Sif S, Jones B, Jackson A, Koipally J, Heller E, et al. Ikaros DNA-binding proteins direct formation of chromatin remodeling complexes in lymphocytes. Immunity. 1999;10(3):345–55.

15. Brown KE, Guest SS, Smale ST, Hahm K, Merkenschlager M, Fisher AG. Association of transcriptionally silent genes with Ikaros complexes at centromeric heterochromatin. Cell. 1997;91(6):845–54.

16. Ge Z, Song EJ, Kawasawa YI, Li J, Dovat S, Song C. WDR5 high expression and its effect on tumorigenesis in leukemia. Oncotarget. 2016;7(25):37740–54.

17. Ge Z, Zhou X, Gu Y, Han Q, Li J, Chen B, et al. Ikaros regulation of the BCL6/BACH2 axis and its clinical relevance in acute lymphoblastic leukemia. Oncotarget. 2017;8(5):8022–34.

18. Ge Z, Gu Y, Xiao L, Han Q, Li J, Chen B, et al. Co-existence of IL7R high and SH2B3 low expression distinguishes a novel high-risk acute lymphoblastic leukemia with Ikaros dysfunction. Oncotarget. 2016;7(29):46014–27.

19. Ge Z, Gu Y, Zhao G, Li J, Chen B, Han Q, et al. High CRLF2 expression associates with IKZF1 dysfunction in adult acute lymphoblastic leukemia without CRLF2 rearrangement. Oncotarget. 2016;7(31):49722–32.

20. Ge Z, Guo X, Li J, Hartman M, Kawasawa YI, Dovat S, et al. Clinical significance of high c-MYC and low MYCBP2 expression and their association with Ikaros dysfunction in adult acute lymphoblastic leukemia. Oncotarget. 2015;6(39):42300–11.

21. Ge Z, Gu Y, Han Q, Zhao G, Li M, Li J, et al. Targeting High Dynamin-2 (DNM2) Expression by Restoring Ikaros Function in Acute Lymphoblastic Leukemia. Sci Rep. 2016;6:38004.

22. Koipally J, Renold A, Kim J, Georgopoulos K. Repression by Ikaros and Aiolos is mediated through histone deacetylase complexes. EMBO J. 1999;18(11):3090–100.

23. Song C, Pan X, Ge Z, Gowda C, Ding Y, Li H, et al. Epigenetic regulation of gene expression by Ikaros, HDAC1 and Casein Kinase II in leukemia. Leukemia. 2016;30(6):1436–40.

24. Ge Z, Song C, Ding Y, Tan BH, Desai D, Sharma A, et al. Dual targeting of MTOR as a novel therapeutic approach for high-risk B-cell acute lymphoblastic leukemia. Leukemia. 2021.

25. Song C, Ge Z, Ding Y, Tan BH, Desai D, Gowda K, et al. IKAROS and CK2 regulate expression of BCL-XL and chemosensitivity inhigh-risk B-cell acute lymphoblastic leukemia. Blood. 2020.

26. Payne JL, Song C, Ding Y, Dhanyamraju PK, Bamme Y, Schramm JW, et al. Regulation of Small GTPase Rab20 by Ikaros in B-Cell Acute Lymphoblastic Leukemia. Int J Mol Sci. 2020;21(5).

27. Ding Y, Zhang B, Payne JL, Song C, Ge Z, Gowda C, et al. Ikaros tumor suppressor function includes induction of active enhancers and super-enhancers along with pioneering activity. Leukemia. 2019;33(11):2720–31.

28. Chen Q, Cui L, Hu Y, Chen Z, Gao Y, Shi Y. Short-term efficacy and safety of biologics and Janus kinase inhibitors for patients with atopic dermatitis: A systematic review and meta-analysis. Heliyon. 2023;9(11):e22014.

29. Brown KE, Baxter J, Graf D, Merkenschlager M, Fisher AG. Dynamic repositioning of genes in the nucleus of lymphocytes preparing for cell division. Mol Cell. 1999;3(2):207–17.

30. Ferreiros-Vidal I, Carroll T, Taylor B, Terry A, Liang Z, Bruno L, et al. Genome-wide identification of Ikaros targets elucidates its contribution to mouse B-cell lineage specification and pre-B-cell differentiation. Blood. 2013;121(10):1769–82.

31. Su RC, Brown KE, Saaber S, Fisher AG, Merkenschlager M, Smale ST. Dynamic assembly of silent chromatin during thymocyte maturation. Nat Genet. 2004;36(5):502–6.

32. Zhang J, Jackson AF, Naito T, Dose M, Seavitt J, Liu F, et al. Harnessing of the nucleosome-remodeling-deacetylase complex controls lymphocyte development and prevents leukemogenesis. Nature immunology. 2012;13(1):86–94.

33. Song C, Gowda C, Pan X, Ding Y, Tong Y, Tan BH, et al. Targeting casein kinase II restores Ikaros tumor suppressor activity and demonstrates therapeutic efficacy in high-risk leukemia. Blood. 2015;126(15):1813–22.

34. Hu Y, Zhang Z, Kashiwagi M, Yoshida T, Joshi I, Jena N, et al. Superenhancer reprogramming drives a B-cell-epithelial transition and high-risk leukemia. Genes Dev. 2016;30(17):1971–90.

35. Kathrein KL, Lorenz R, Innes AM, Griffiths E, Winandy S. Ikaros induces quiescence and T-cell differentiation in a leukemia cell line. Mol Cell Biol. 2005;25(5):1645–54.

36. Wang Z, Zang C, Rosenfeld JA, Schones DE, Barski A, Cuddapah S, et al. Combinatorial patterns of histone acetylations and methylations in the human genome. Nat Genet. 2008;40(7):897–903.

37. Fujiwara T, O’Geen H, Keles S, Blahnik K, Linnemann AK, Kang YA, et al. Discovering hematopoietic mechanisms through genome-wide analysis of GATA factor chromatin occupancy. Mol Cell. 2009;36(4):667–81.

38. Kirmizis A, Bartley SM, Kuzmichev A, Margueron R, Reinberg D, Green R, et al. Silencing of human polycomb target genes is associated with methylation of histone H3 Lys 27. Genes Dev. 2004;18(13):1592–605.

39. Mochizuki-Kashio M, Mishima Y, Miyagi S, Negishi M, Saraya A, Konuma T, et al. Dependency on the polycomb gene Ezh2 distinguishes fetal from adult hematopoietic stem cells. Blood. 2011;118(25):6553–61.

40. Oravecz A, Apostolov A, Polak K, Jost B, Le Gras S, Chan S, et al. Ikaros mediates gene silencing in T cells through Polycomb repressive complex 2. Nat Commun. 2015;6:8823.

41. Koipally J, Kim J, Jones B, Jackson A, Avitahl N, Winandy S, et al. Ikaros chromatin remodeling complexes in the control of differentiation of the hemo-lymphoid system. Cold Spring Harb Symp Quant Biol. 1999;64:79–86.

42. Cobb BS, Morales-Alcelay S, Kleiger G, Brown KE, Fisher AG, Smale ST. Targeting of Ikaros to pericentromeric heterochromatin by direct DNA binding. Genes and Development. 2000;14:2146–60.

43. Crispatzu G, Rehimi R, Pachano T, Bleckwehl T, Cruz-Molina S, Xiao C, et al. The chromatin, topological and regulatory properties of pluripotency-associated poised enhancers are conserved in vivo. Nat Commun. 2021;12(1):4344.

44. Juric I, Yu M, Abnousi A, Raviram R, Fang R, Zhao Y, et al. MAPS: Model-based analysis of long-range chromatin interactions from PLAC-seq and HiChIP experiments. PLoS Comput Biol. 2019;15(4):e1006982.

45. Madani Tonekaboni SA, Haibe-Kains B, Lupien M. Large organized chromatin lysine domains help distinguish primitive from differentiated cell populations. Nat Commun. 2021;12(1):499.

46. Zhao D, Zhang L, Zhang M, Xia B, Lv J, Gao X, et al. Broad genic repression domains signify enhanced silencing of oncogenes. Nat Commun. 2020;11(1):5560.

47. Hasegawa N, Oshima M, Sashida G, Matsui H, Koide S, Saraya A, et al. Impact of combinatorial dysfunctions of Tet2 and Ezh2 on the epigenome in the pathogenesis of myelodysplastic syndrome. Leukemia. 2017;31(4):861–71.

48. Iwama A. Polycomb repressive complexes in hematological malignancies. Blood. 2017;130(1):23–9.

49. Sashida G, Harada H, Matsui H, Oshima M, Yui M, Harada Y, et al. Ezh2 loss promotes development of myelodysplastic syndrome but attenuates its predisposition to leukaemic transformation. Nat Commun. 2014;5:4177.

50. Sashida G, Iwama A. Multifaceted role of the polycomb-group gene EZH2 in hematological malignancies. Int J Hematol. 2017;105(1):23–30.

51. Sashida G, Wang C, Tomioka T, Oshima M, Aoyama K, Kanai A, et al. The loss of Ezh2 drives the pathogenesis of myelofibrosis and sensitizes tumor-initiating cells to bromodomain inhibition. J Exp Med. 2016;213(8):1459–77.

52. van Dijk AD, Hoff FW, Qiu YH, Chandra J, Jabbour E, de Bont E, et al. Loss of H3K27 methylation identifies poor outcomes in adult-onset acute leukemia. Clin Epigenetics. 2021;13(1):21.

53. Dunham I, Kundaje A, Aldred SF, Collins PJ, Davis CA, Doyle F, et al. An integrated encyclopedia of DNA elements in the human genome. Nature. 2012;489(7414):57–74.

54. Wang C, Oshima M, Sato D, Matsui H, Kubota S, Aoyama K, et al. Ezh2 loss propagates hypermethylation at T cell differentiation-regulating genes to promote leukemic transformation. J Clin Invest. 2018;128(9):3872–86.

55. Li B, Chng WJ. EZH2 abnormalities in lymphoid malignancies: underlying mechanisms and therapeutic implications. J Hematol Oncol. 2019;12(1):118.

56. Ntziachristos P, Tsirigos A, Van Vlierberghe P, Nedjic J, Trimarchi T, Flaherty MS, et al. Genetic inactivation of the polycomb repressive complex 2 in T cell acute lymphoblastic leukemia. Nature medicine. 2012;18(2):298–301.

57. Kuzmichev A, Nishioka K, Erdjument-Bromage H, Tempst P, Reinberg D. Histone methyltransferase activity associated with a human multiprotein complex containing the Enhancer of Zeste protein. Genes Dev. 2002;16(22):2893–905.

## References

1. Langmead B, Salzberg SL. Fast gapped-read alignment with Bowtie 2. Nat Methods. 2012;9(4):357–9.

2. Li H.* HB, Wysoker A., Fennell T., Ruan J., Homer N., Marth G., Abecasis G., Durbin R. and 1000 Genome Project Data Processing Subgroup The Sequence alignment/map (SAM) format and SAMtools. Bioinformatics. 2009(25):2078–9.

3. al Ze. Model-based Analysis of ChIP-Seq (MACS). Genome Biol. 2008;9(9):137.

4. al Le. ChIP-seq guidelines and practices of the ENCODE and modENCODE consortia. Genome Res. 2012(22):1813–31.

5. Madani Tonekaboni SA, Haibe-Kains B, Lupien M. Large organized chromatin lysine domains help distinguish primitive from differentiated cell populations. Nat Commun. 2021;12(1):499.

6. Zhao D, Zhang L, Zhang M, Xia B, Lv J, Gao X, et al. Broad genic repression domains signify enhanced silencing of oncogenes. Nat Commun. 2020;11(1):5560.

7. Martin, M. Cutadapt Removes Adapter Sequences From High-Throughput Sequencing Reads.

8. Liao Y, Smyth GK, Shi W. featureCounts: an efficient general purpose program for assigning sequence reads to genomic features. Bioinformatics. 2014;30(7):923–30.

9. Durinck S SP, Birney E, Huber W Mapping identifiers for the integration of genomic datasets with the R/Bioconductor package biomaRt. Nature Protocols. 2009;4(4):1184–91.

10. He B, Chen C, Teng L, Tan K. Global view of enhancer-promoter interactome in human cells. Proc Natl Acad Sci U S A. 2014;111(21):E2191–9.

11. Ramirez F, Ryan DP, Gruning B, Bhardwaj V, Kilpert F, Richter AS, et al. deepTools2: a next generation web server for deep-sequencing data analysis. Nucleic Acids Res. 2016;44(W1):W160–5.

12. Timothy L. Bailey MB, Fabian A. Buske, Martin Frith, Charles E. Grant, Luca Clementi, Jingyuan Ren, Wilfred W. Li, William S. Noble. MEME SUITE: tools for motif discovery and searching. Nucleic Acids Research. 2009;37:W202–W8.

13. Huang DW SB, Lempicki RA. Systematic and integrative analysis of large gene lists using DAVID Bioinformatics Resources. Nature Protocols. 2009;4(1):44–57.

14. Cory Y, McLean DB, Michael Hiller, Shoa L Clarke, Bruce T Schaar, Craig B Lowe, Aaron M Wenger, and Gill Bejerano. GREAT improves functional interpretation of cis-regulatory regions. Nat Biotechnol. 2010;28(5):495–501.

